# Mitotic clustering of pulverized chromosomes from micronuclei

**DOI:** 10.1101/2022.07.19.500697

**Authors:** Yu-Fen Lin, Jose Espejo Valle-Inclan, Alice Mazzagatti, Qing Hu, Elizabeth G. Maurais, Alison Guyer, Jacob T. Sanders, Justin Engel, Giaochau Nguyen, Daniel Bronder, Samuel F. Bakhoum, Isidro Cortés-Ciriano, Peter Ly

## Abstract

Complex genome rearrangements can be generated by the catastrophic shattering of mis-segregated chromosomes trapped within micronuclei through a process known as chromothripsis. Since each chromosome harbors a single centromere, how acentric fragments derived from shattered chromosomes are inherited between daughter cells during mitosis remains unknown. Here we tracked micronucleated chromosomes by live-cell imaging and show that acentric fragments cluster in close spatial proximity throughout mitosis for biased partitioning to a single daughter cell. Mechanistically, the CIP2A-TOPB1 complex prematurely associates with DNA lesions within ruptured micronuclei during interphase, which poises chromosome fragments for clustering upon mitotic entry. Inactivation of CIP2A or TOPBP1 caused pulverized chromosomes to untether and disperse throughout the mitotic cell, consequently resulting in the mis-accumulation of DNA fragments in the cytoplasm. The inheritance of shattered chromosomes by a single daughter cell suggests that micronucleation can drive complex rearrangements that lack the DNA copy number oscillations characteristic of canonical chromothripsis. Comprehensive analysis of pan-cancer whole-genome sequencing data revealed clusters of DNA copy number-neutral rearrangements – termed balanced chromothripsis – across diverse cancer types resulting in the acquisition of known driver events. Thus, distinct patterns of chromothripsis can be explained by the spatial mitotic clustering of pulverized chromosomes from micronuclei.

---

Chromothripsis drives complex genome rearrangements through the catastrophic pulverization of individual chromosomes^1^. In cancer genomes, these rearrangements are accompanied by a characteristic DNA copy number pattern that oscillates between two states, representing the retention and loss of fragments along the shattered chromosome^1-4^. Chromothripsis can be initiated by mitotic cell division errors resulting in the encapsulation of mis-segregated chromosomes into abnormal nuclear structures called micronuclei^5-8^, which acquire extensive DNA damage in interphase upon rupture of its nuclear membrane^9-11^. Following entry into mitosis, damaged chromosomes within micronuclei pulverize into dozens of microscopic fragments^12,13^.

The stochastic inheritance of chromosome fragments by both newly formed daughter cells could in part contribute to the alternating DNA copy number states characteristic of chromothripsis^5,14^. Sequencing of daughter cell pairs derived from micronucleated mother cells showed that complex rearrangements are indeed a common outcome of micronucleus formation^5^. However, in most cases, the patterns of chromothripsis differed from those in cancer genomes as the rearrangements were largely restricted to a single daughter cell and lacked the canonical oscillations in DNA copy number states^5^. Moreover, germline chromothripsis events reported in congenital disorders typically generate complex yet balanced rearrangements, indicative of minimal DNA loss^15,16^. These studies implicate a potential mechanism suppressing the loss of genetic material following chromosome pulverization, although how distinct patterns of rearrangements arise in cancers and germline disorders remain unclear. In metazoans, the maintenance of a single centromere per chromosome is critical for establishing bipolar attachments to the mitotic spindle and achieving high-fidelity genome segregation^17^. However, most fragments derived from micronuclei are acentric and cannot directly bind to spindle microtubules^13^. Several models have been proposed to promote acentric chromosome segregation, including proteins that function as DNA tethers^18^, the hitchhiking of extrachromosomal DNA (ecDNA) with linear chromosomes^19^, tethering of viral episomes to host chromosomes^20,21^, and controlled delays in nuclear envelope reassembly or cytokinesis^22,23^. Despite this knowledge, we have an incomplete understanding of how acentric fragments from micronuclei are inherited by daughter cells during mitosis, an important step for its reassembly upon reincorporation into the interphase nucleus.

We previously developed a human cell-based system to induce the formation of micronuclei harboring the Y chromosome in genome-engineered, chromosomally stable DLD-1 cells^6,13^. Following exposure to doxycycline and auxin (DOX/IAA), the centromere-specific histone H3 variant CENP-A is replaced with a CENP-A mutant that functionally inactivates the Y centromere without disrupting autosome or X chromosome segregation^13,24,25^. This system recapitulates the stepwise events of chromothripsis, including the pulverization of mis-segregated chromosomes in micronuclei^13^ that result in complex rearrangements^6^. Building upon this approach to monitor the partitioning of acentric chromosome fragments, here we show that pulverized chromosomes from micronuclei spatially cluster throughout mitosis and identify the CIP2A-TOPBP1 complex as an essential regulator of this process. Mitotic clustering drives the unequal inheritance of acentric fragments by a single daughter cell, providing an explanation for the origins of distinct patterns of chromothripsis found across diverse cancer types and congenital disorders.

## Results

### Pulverized chromosomes cluster throughout mitosis for unequal segregation

To visualize the behavior of micronucleated chromosomes during mitosis by live-cell imaging, we labeled the Y chromosome in DLD-1 cells using a nuclease-dead Cas9 (dCas9) fused to a SunTag scaffold (dCas9-SunTag, see **Methods**), which can recruit 10-24 copies of superfolder green fluorescent protein (sfGFP) fused to a single-chain variable fragment^26^. Since the Y chromosome q-arm is comprised of a ∼30 megabase (Mb) repeat array termed DYZ1^27^, we reasoned that a single sgRNA targeting DYZ1 could tile multiple dCas9-SunTag copies across half of the Y chromosome for labeling with sfGFP (**Fig. 1a**). To identify optimal CRISPR target sequences, we first generated stable cell lines encoding individual candidate sgRNAs, leading to the identification of a DYZ1 sgRNA that produced a single, nuclear sfGFP signal (**Extended Data Fig. 1a-b**). To reduce heterogeneity in expression levels, clones derived from the parental population expressing sfGFP under the control of various promoters or harboring different SunTag scaffold lengths were then screened for optimal, homogenous sfGFP levels (**Extended Data Fig. 1c-d**). As expected, dCas9-SunTag strongly co-localized with DNA fluorescence *in situ* hybridization (FISH) probes targeting the Y chromosome in interphase nuclei and on metaphase spreads (**Extended Data Fig. 1e-g**).

**Figure 1.**
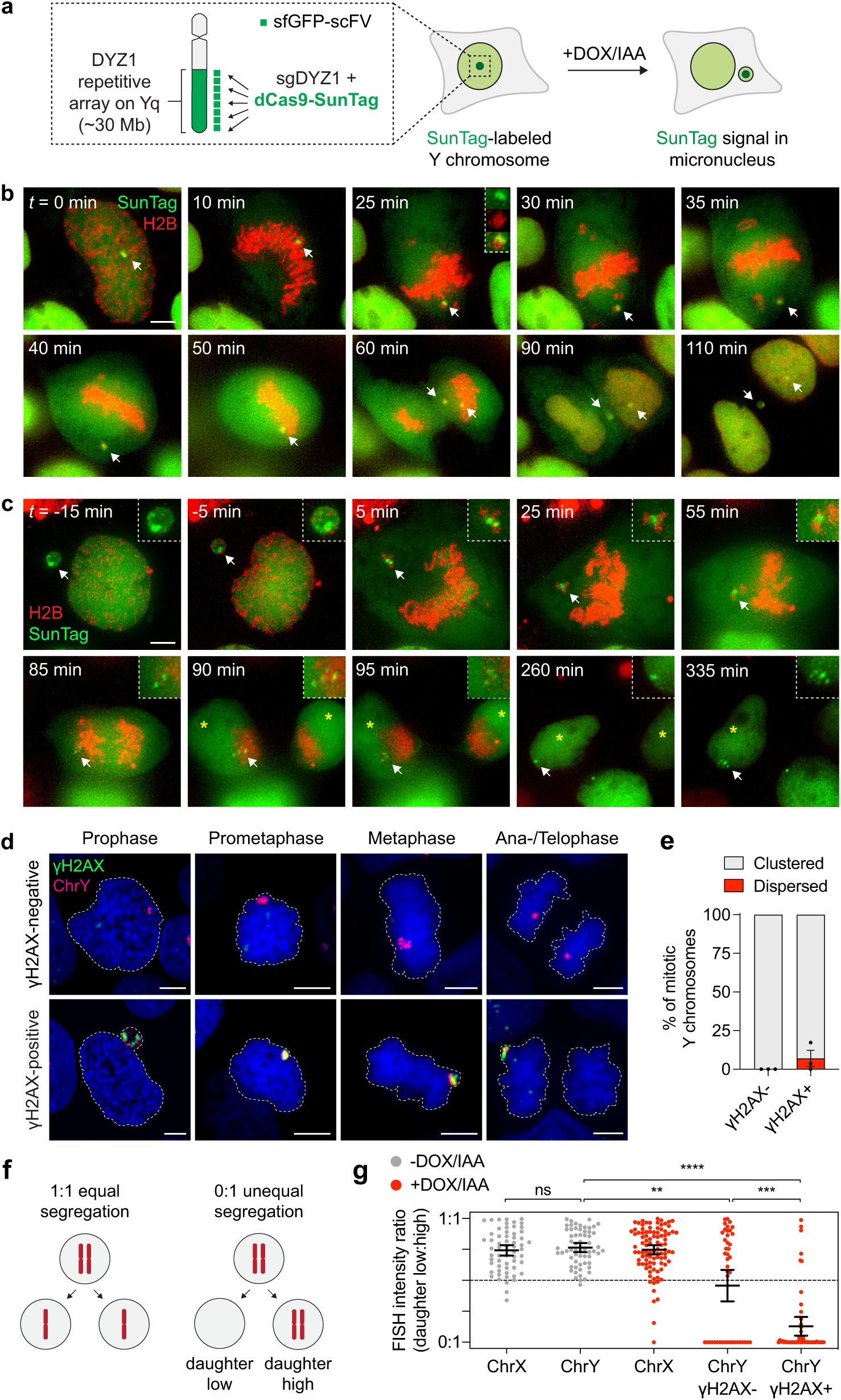
Pulverized chromosomes from micronuclei remain clustered throughout mitosis for biased inheritance by a single daughter cell. **a)** Schematic of chromosome-labeling strategy using a dCas9-SunTag fusion to target superfolder green fluorescent protein fused to scFV (sfGFP-scFV) to the DYZ1 satellite array located on the Y chromosome q-arm. Treatment with doxycycline and auxin (DOX/IAA) triggers Y centromere inactivation and Y chromosome mis-segregation into micronuclei. **b)** Time-lapse images of a dCas9-SunTag-labeled Y chromosome (white arrow) mis-segregating into a micronucleus during mitosis following 2d DOX/IAA induction. Scale bar, 5 μm. **c)** Time-lapse images of a dCas9-SunTag-labeled Y chromosome in a micronucleus (white arrow) clustering throughout mitosis with uneven distribution of the SunTag signal between daughter cells. Yellow asterisks denote the two newly formed daughter cells. Representative example shown from *n* = 13 events obtained from independent experiments. In (**b**) and (**c**), chromatin is labeled with H2B-mCherry. Scale bar, 5 μm. **d)** Confirmation of micronuclear chromosome clustering in fixed DLD-1 cells at different stages of mitosis with chromosome-paint DNA fluorescence *in situ* hybridization (FISH) probes. Staining for γH2AX is used to identify pulverized chromosomes from micronuclei harboring extensive DNA double-strand breaks (DSBs). Blue, 4′,6-diamidino-2-phenylindole (DAPI) stain for DNA. Scale bar, 5 μm. **e)** Quantification of fragment clustering with and without γH2AX from prometaphase to metaphase. Data represent mean ± SEM of *n* = 3 independent experiments; γH2AX-negative = 595, γH2AX-positive = 230 cells. **f)** Schematic of chromosome distribution in a 1:1 or 1:0 segregation ratio between daughter cells. **g)** Pulverized chromosomes labeled with γH2AX are inherited by a single daughter cell. Data represent mean ± 95 confidence intervals; not significant (ns), *P* > 0.05, \*\**P* = 0.0026, \*\*\**P* = 0.0006, \*\*\*\**P* ≤ 0.0001 by non-parametric Kruskal-Wallis test with multiple comparisons; left to right: *n* = 61, 61, 95, 44, and 51 pairs of daughter cells pooled from 3 independent experiments. Representative images are shown in **Extended Data Fig. 2f**.

Using this chromosome-labeling strategy, we tested whether Y centromere inactivation could trigger the mis-segregation of the dCas9-SunTag-labeled Y chromosome into micronuclei. Treatment with DOX/IAA efficiently relocated dCas9-SunTag signals from the nucleus to micronuclei (**Extended Data Fig. 1h-i**). Time-lapse microscopy confirmed frequent mis-segregation of the Y chromosome into micronuclei during mitosis (**Fig. 1b**). In the example shown, sister chromatid disjunction can be observed during anaphase by the resolution of a single SunTag puncta into two discrete signals, one of which mis-segregates into a micronucleus (**Fig. 1b**). To examine the fate of Y chromosome fragments in mitosis, cells with dCas9-SunTag-labeled micronuclei were first arrested in G2 phase using a CDK1 inhibitor. Most micronuclei had reduced background levels of diffused sfGFP, indicating that a proportion of observed events represented ruptured micronuclei with defective nucleocytoplasmic compartmentalization (**Extended Data Fig. 1j**) – a hallmark feature of micronuclei^9^. Following release from G2, live-cell imaging revealed that micronuclear fragments unexpectedly remained clustered as a discrete dCas9-SunTag signal throughout mitosis in all events captured (*n* = 13, **Fig. 1c**). Notably, mitotic clustering resulted in the biased partitioning of most fragments into a single daughter nucleus (**Fig. 1c**), highlighting an unidentified mechanism that tethers chromosome fragments for biased partitioning as a collective unit.

To confirm these findings from live cells, we fixed and stained intact mitotic cells for γH2AX as a surrogate for micronuclear fragments harboring DNA double-strand breaks (DSBs). Following DOX/IAA treatment, nearly one-third of mitotic cells (31%, *n* = 541 cells) exhibited γH2AX signals that were specific for the Y chromosome. In agreement with live-cell dCas9-SunTag imaging, the majority (∼94%) of γH2AX-positive Y chromosome fragments were confined to a small, discrete region (**Fig. 1d-e**). Fragment clustering was observed throughout the entire duration of mitosis, indicating that they do not resolve at a specific mitotic stage, and indeed, the majority of signals were inherited by a single daughter cell following anaphase onset (**Fig. 1d**). Analyses of interphase nuclei at the corresponding time point revealed clusters of damaged chromosome fragments (**Extended Data Fig. 2a-c**) with expanded fluorescence signal (**Extended Data Fig. 2d-e**) that resembled subnuclear territories termed micronuclei bodies^28^. The nuclear envelope of micronuclei undergoes efficient disassembly during mitosis^6^, suggesting that fragment clustering is not caused by failures in micronuclear envelope breakdown. Altogether, pulverized fragments from micronuclei have an intrinsic ability to cluster throughout mitotic cell division.

We next explored the unequal inheritance patterns of fragmented chromosomes during mitosis. To do so, we quantified the intensity of Y chromosome FISH probes between pairs of newly formed daughter cells (**Fig. 1f**). In unperturbed conditions, the Y chromosome and a control X chromosome segregated at an expected 1:1 ratio (**Fig. 1f**, grey points). Following Y centromere inactivation, 57% of intact Y chromosomes (γH2AX-negative, *n* = 44 daughter pairs) segregated normally (ratio > 0.5:1), whereas 43% underwent whole-chromosome segregation errors resulting in complete loss of the Y in one daughter cell (ratio = 0:1, **Fig. 1f**, red points). In contrast, most (88%) pulverized Y chromosomes (γH2AX-positive, *n* = 51 daughter pairs) were frequently inherited at an unequal ratio approaching 0:1, consistent with the segregation of most fragments into a single daughter nucleus (**Fig. 1f**). These data demonstrate that mitotic clustering of acentric fragments originating from micronuclei promotes the biased partitioning of pulverized chromosomes to one daughter cell.

### CIP2A-TOPBP1 is essential for mitotic fragment clustering

Several candidates of the DNA damage response have been implicated in facilitating the segregation of acentric chromosomes and/or suppressing micronuclei formation, including DNA polymerase theta (encoded by *POLQ*) and the MRE11-RAD50-NBS1 (MRN) complex in flies^29^, as well as the CIP2A-TOPBP1 complex in mammalian cells^30-32^. To determine whether these DNA damage response components are involved in fragment clustering, we assessed mitotic chromosome clustering in CRISPR/Cas9-edited *POLQ*^−/−^, *NBN*^−/−^, and *CIP2A*^−/−^ DLD-1 clones (**Fig. 2a**). Whereas DNA polymerase theta and NBS1 knockout (KO) cells were similar to wildtype (WT) controls, cells lacking CIP2A exhibited a striking dispersion of Y chromosome fragments (**Fig. 2b**) across independent clones generated with distinct sgRNAs targeting exon 1 (indicated as sg3 and sg4, **Extended Data Fig. 3a**).

**Figure 2.**
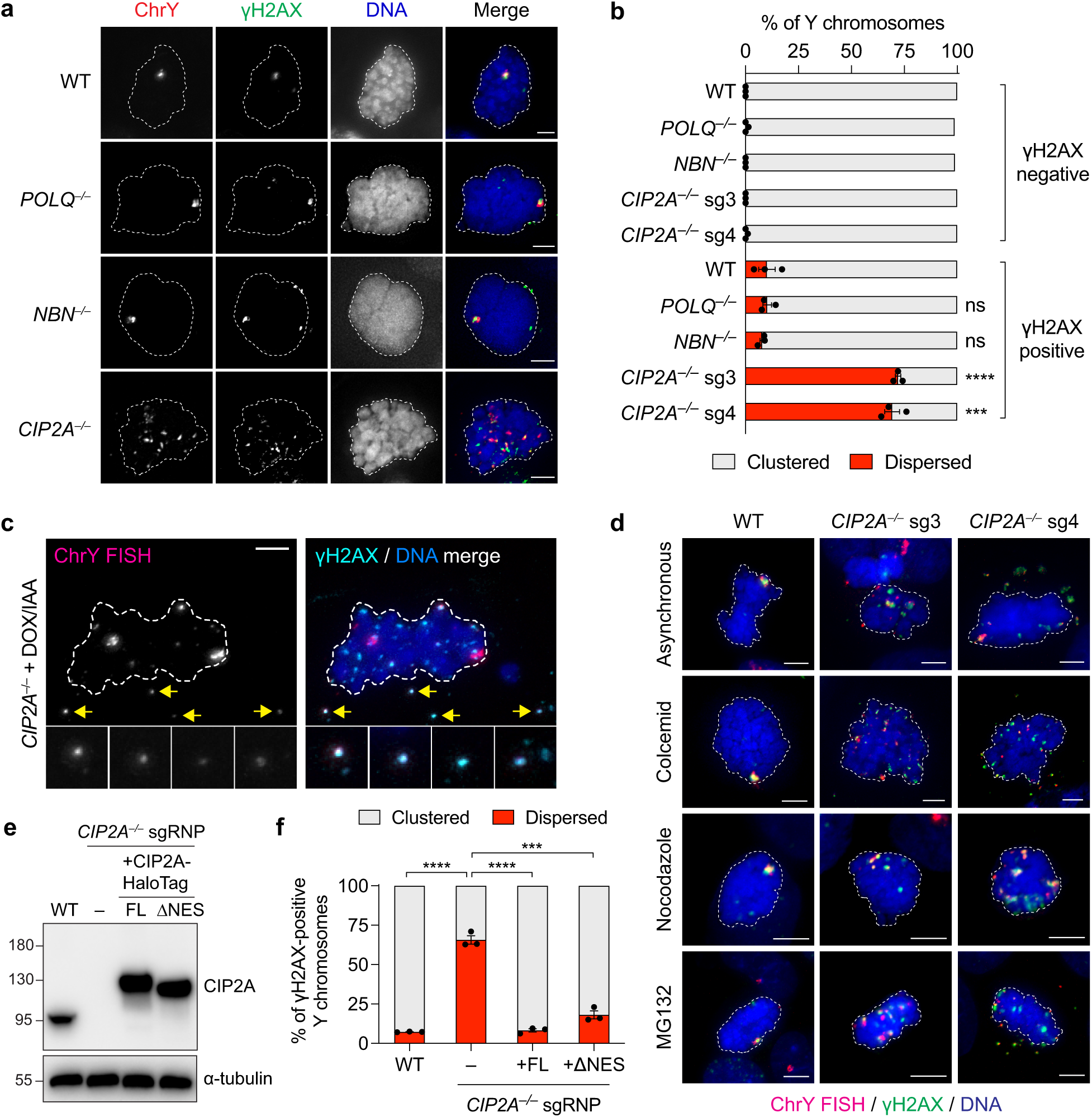
Inactivation of CIP2A disrupts clustering and disperses pulverized chromosomes throughout the mitotic cytoplasm. **a)** Images of DOX/IAA-treated mitotic DLD-1 cells with the indicated biallelic gene deletions with γH2AX-positive Y chromosomes. sg3 and sg4 denote distinct sgRNAs targeting exon 1 of CIP2A. Scale bar, 5 μm. **b)** Frequency of fragment clustering from γH2AX-negative and γH2AX-positive Y chromosomes from (**a**). Data represent mean ± SEM; not significant (ns), *P* > 0.05, \*\*\**P* = 0.0003, \*\*\*\**P* ≤ 0.0001 by two-tailed t-test compared to WT controls; *n* = 3 independent experiments; top to bottom: 657, 455, 352, 526, 383, 238, 165, 175, 179, and 149 cells. **c)** Example of mitotic CIP2A knockout (KO, sg3) cell with γH2AX-positive Y chromosome fragments that are displaced from the main chromosome mass (yellow arrows). Scale bar, 5 μm. **d)** Images of mitotic CIP2A KO cells exhibiting fragment dispersion in randomly cycling cells and cells arrested with the indicated mitotic inhibitors. Scale bar, 5 μm. **e)** Western blot confirmation of ectopic CIP2A-HaloTag complementation in CIP2A KO cells generated by a frameshift deletion in exon 3 induced by Cas9 ribonucleoprotein (RNP) delivery. FL, full length; NES, nuclear export signal. **f)** Frequency of fragment clustering from γH2AX-positive Y chromosomes from CIP2A rescue cells. Data represent mean ± SEM; \*\*\*\**P* ≤ 0.0001, \*\*\**P* = 0.0002 by two-tailed t-test compared to CIP2A KO cells; *n* = 3 independent experiments; WT = 137, no rescue = 200, FL rescue = 139, delta NES rescue = 115 cells.

CIP2A KO cells proliferated at a slightly reduced rate (**Extended Data Fig. 3b**) but exhibited a normal cell cycle distribution profile (**Extended Data Fig. 3c**). As predicted, treatment with DOX/IAA triggered micronucleation and Y chromosome fragmentation at a frequency comparable to WT cells (**Extended Data Fig. 3d-e**), suggesting that CIP2A is not directly involved in driving whole-chromosome segregation errors or DNA damage formation within micronuclei. However, mitotic CIP2A KO cells harbored Y chromosome fragments co-localizing with γH2AX that were noticeably displaced from the primary genomic mass (**Fig. 2c**). Up to 45 distinct fragments were detected by microscopy in CIP2A KO cells, whereas WT cells infrequently (13.6%, *n* = 214 cells) contained >5 dispersed fragments (**Extended Data Fig. 4a**). Fragment dispersion was evident in unsynchronized mitotic cells and those arrested with inhibitors that interfered with microtubule polymerization or the spindle assembly checkpoint (**Fig. 2d**). Complementation of CIP2A KO cells (generated by a frameshift deletion in exon 3) with full-length CIP2A fused to a HaloTag (CIP2A-HaloTag) fully rescued mitotic fragment clustering (**Fig. 2e-f**). Depletion of either CIP2A or its interacting partner TOPBP1^30,32-34^ in WT cells by independent small interfering RNAs (siRNAs) (**Extended Data Fig. 4b**) was sufficient to disrupt mitotic clustering and disperse chromosome fragments (**Extended Data Fig. c-d**). Acute loss of CIP2A also increased the number of detectable dCas9-SunTag-labeled Y chromosome fragments in live cells (**Extended Data Fig. 4e-f**).

To determine the spatial arrangement of micronuclear chromosome fragments, we analyzed metaphase spreads prepared from mitotic DLD-1 cells swollen by hypotonic treatment. Following the induction of micronucleation and chromosome fragmentation, DNA FISH using Y chromosome painting probes revealed different degrees of fragment spreading, ranging from those that remained in close proximity to those that were scattered throughout the metaphase spread area (**Extended Data Fig. 5a-c**). Notably, both CIP2A KO cells and WT cells depleted of CIP2A or TOPBP1 displayed a higher degree of metaphase chromosome fragment dispersion (**Extended Data Fig. 5d-e**). Altogether, these efforts identify the CIP2A-TOPBP1 complex as an essential mitotic regulator involved in the spatial clustering of pulverized chromosomes in micronuclei.

### Micronuclear CIP2A-TOPBP1 poises acentric fragments for clustering

To determine whether CIP2A associates with micronuclear fragments prior to mitotic entry, we assessed the interphase localization of CIP2A, which contains a nuclear export signal (NES) that drives its compartmentalization within the cytoplasm^32^. Consistent with this, CIP2A was rarely observed within intact micronuclei (**Fig. 3a, Extended Data Fig. 6a**). However, using γH2AX as a marker of micronuclear envelope rupture^9^, two distinct patterns of CIP2A emerged. First, diffused CIP2A matching the intensity of the cytoplasmic pool of CIP2A was observed in one-third (∼38%) of ruptured micronuclei (**Fig. 3a-b, Extended Data Fig. 6a**), which is likely caused by defects in nucleocytoplasmic compartmentalization. Second, CIP2A appeared as robust puncta, which were less frequent (∼7%) but displayed a strong association with micronuclear DNA damage (**Fig. 3a-b, Extended Data Fig. 6a**). These patterns were confirmed using the accumulation of cGAS-GFP as an alternative marker for ruptured micronuclei^35,36^ (**Extended Data Fig. 7a**,**c**). Thus, whereas CIP2A does not associate with interphase DSBs in the nucleus^30,32,37^, we propose that cytoplasmic CIP2A diffuses into ruptured micronuclei where it prematurely engages with micronuclear DSBs that accumulate throughout interphase. In agreement, a mutant CIP2A rescue lacking its NES was sufficient to restore fragment clustering in CIP2A KO cells (**Fig. 2f**), suggesting that the normal cytoplasmic localization of CIP2A is dispensable for mitotic clustering following micronuclear envelope rupture.

**Figure 3.**
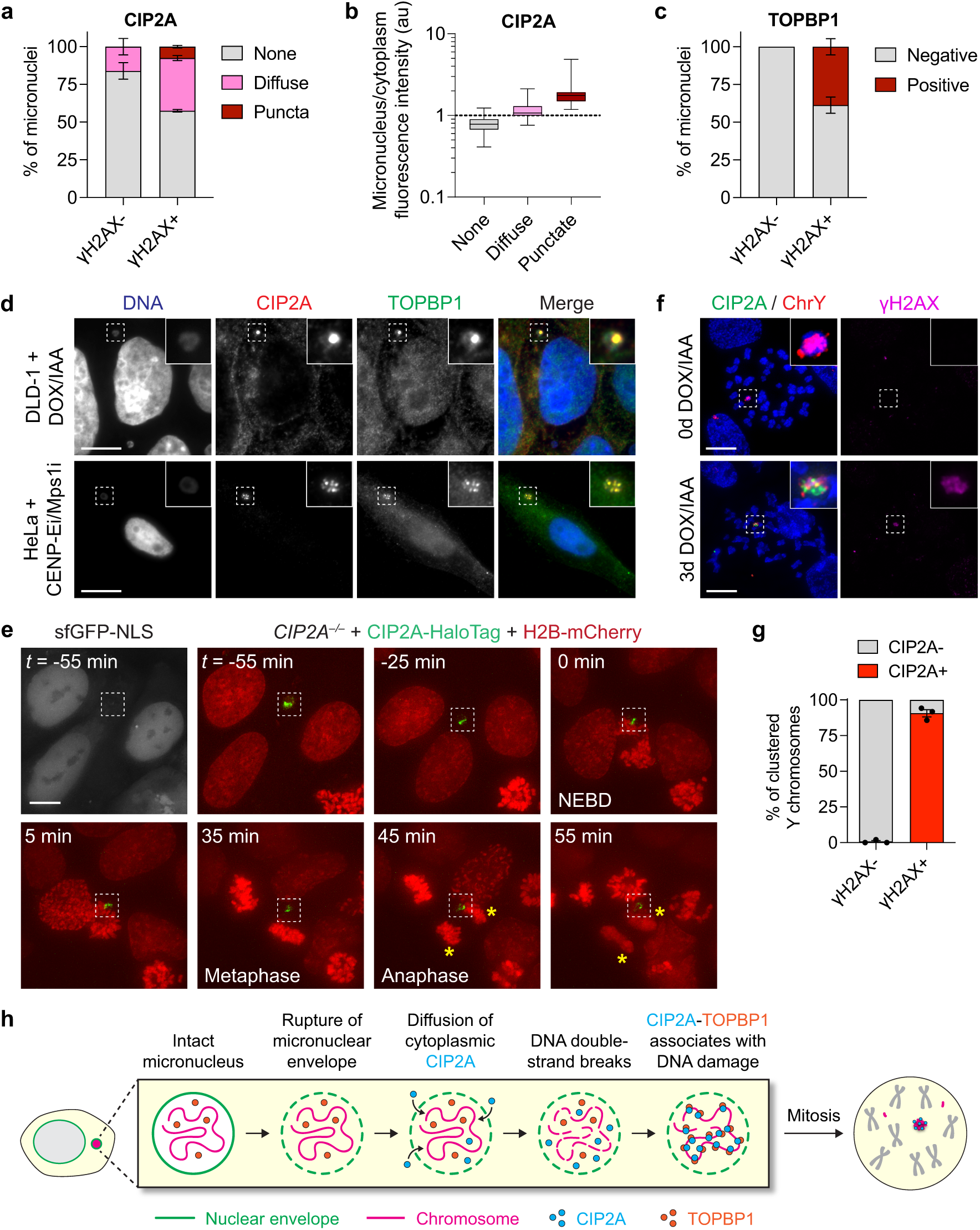
Recruitment of CIP2A-TOPBP1 to ruptured micronuclei poises acentric fragments for clustering upon mitotic entry. **a)** Frequency of CIP2A localization patterns in intact (γH2AX-negative) and ruptured (γH2AX-positive) micronuclei of DLD-1 cells. Data represent mean ± SEM; *n* = 3 independent experiments; γH2AX-negative = 218, γH2AX-positive = 99 micronuclei. See **Extended Data Fig. 7a** for complete set of images. **b)** Intensity measurements of the indicated CIP2A localization patterns in micronuclei compared to the cytoplasm. Box plot represents interquartile range with min-max; none: *n* = 236, diffuse: *n* = 74, puncta: *n* = 7 micronuclei from 3 independent experiments. au, arbitrary units. **c)** Frequency of TOPBP1 localization patterns in intact (γH2AX-negative) and ruptured (γH2AX-positive) micronuclei. Data represent mean ± SEM; *n* = 3 independent experiments; γH2AX-negative = 323, γH2AX-positive = 120 micronuclei. See **Extended Data Fig. 6b** for complete set of images. **d)** Co-localization of CIP2A and TOPBP1 puncta in micronuclei of DLD-1 cells with Y chromosome micronuclei (left) and HeLa cells with micronuclei harboring random chromosomes (right). Scale bar, 10 μm. See **Extended Data Fig. 7e** and **8f** for complete set of images. **e)** Time-lapse example of interphase CIP2A-HaloTag signal in a ruptured micronucleus (lacking sfGFP-NLS) upon mitotic entry and throughout the completion of mitosis. Yellow asterisks denote the two newly formed daughter cells. NEBD, nuclear envelope breakdown. Scale bar, 5 μm. **f)** Examples of mitotic chromosomes showing a highly specific association between CIP2A with clusters of γH2AX-positive Y chromosome fragments (+DOX/IAA) but not γH2AX-negative Y chromosomes (-DOX/IAA). Scale bar, 10 μm. **g)** Quantification of CIP2A localization on Y chromosomes with and without γH2AX from (**f**). Data represent mean ± SEM; *n* = 3 independent experiments; γH2AX-negative = 611, γH2AX-positive = 200 chromosome spreads. **h)** Schematic depicting the stepwise series of events resulting in the premature engagement of CIP2A-TOPBP1 with DNA damage in ruptured micronuclei during interphase.

TOPBP1 is normally nuclear and sequestered away from cytoplasmic CIP2A^30,32^. Intense TOPBP1 puncta were visible exclusively within ruptured, but not intact, micronuclei (**Fig. 3c, Extended Data Fig. 6b, Extended Data Fig. 7b**,**d**). Notably, CIP2A and TOPBP1 formed highly co-localized puncta within micronuclei in DLD-1 cells (**Fig. 3d, Extended Data Fig. 7e-f**). These interphase associations between CIP2A-TOPBP1 were further confirmed in HeLa S3 cells harboring micronuclei induced by transient co-inhibition of the CENP-E motor protein and MPS1 mitotic kinase (**Fig. 3d, Extended Data Fig. 8**). Thus, cytoplasmic CIP2A and nuclear TOPBP1 prematurely associate with DSBs following the loss of nucleocytoplasmic compartmentalization within ruptured micronuclei during interphase. To visualize this process in live cells, we performed time-lapse imaging of CIP2A KO cells reconstituted with CIP2A-HaloTag (**Fig. 2e**), confirming the interphase puncta localization of CIP2A within ruptured micronuclei, as determined by the absence of sfGFP fused to a nuclear localization signal (**Fig. 3e**). Importantly, as the nuclear envelope disassembled upon mitotic entry, the micronucleated CIP2A-HaloTag puncta remained coalesced until the completion of mitosis, resulting in its partitioning exclusively to a single daughter cell (**Fig. 3e**).

We next examined the localization of CIP2A and TOPBP1 on chromosome fragments during mitosis. Consistent with prior reports^30,32^, CIP2A formed small, co-localized foci with spontaneous mitotic DNA lesions in unperturbed cells (**Extended Data Fig. 9a**, top panel). Following DOX/IAA induction, a highly specific association between large CIP2A puncta with clusters of γH2AX-positive Y chromosome fragments were observed in both mitotic cells (**Extended Data Fig. 9a**, bottom panel) and chromosomes (**Fig. 3f-g**). By contrast, mitotic CIP2A puncta were undetectable in control cells with intact chromosomes (**Fig. 3f-g**). TOPBP1 was similarly recruited to clustered mitotic chromosome fragments in WT cells (**Extended Data Fig. 9b-c**) but not CIP2A KO cells (**Extended Data Fig. 9c-d**), suggesting that TOPBP1 function during mitosis is dependent on CIP2A, as shown for spontaneous mitotic DNA damage^32^. Altogether, we propose that CIP2A-TOPBP1 bound to micronuclear DNA lesions poises acentric fragments for clustering immediately upon mitotic entry, which subsequently tethers pulverized chromosomes in spatial proximity throughout mitosis (**Fig. 3h**).

To determine whether CIP2A-TOPBP1 interacts with acentric chromosomes in the absence of DNA damage, we examined the mitotic localization of CIP2A and TOPBP1 on acentric ecDNA elements, which can be produced by chromothripsis^5,6,38^ but lack DSB ends due to their circularized nature. To do so, we used PC3 cells, which harbor abundant ecDNAs in the form of double minute chromosomes that are visible on metaphase spreads, alongside ecDNA-negative HeLa S3 cells as a control (**Extended Data Fig. 10a**). In unperturbed mitotic PC3 and HeLa S3 cells, both CIP2A and TOPBP1 foci were not visible except for those that co-localized with spontaneous DSBs (**Extended Data Fig. 10b-c**). In contrast, DSBs induced by ionizing radiation stimulated the appearance of both mitotic CIP2A and TOPBP1 foci (**Extended Data Fig. 10b-c**). Thus, the CIP2A-TOPBP1 complex is not normally recruited to ecDNAs during mitosis, indicating that its association with acentric chromosome fragments following chromothripsis requires the presence of DNA damage.

### Mitotic clustering suppresses the accumulation of cytoplasmic DNA

In the absence of CIP2A, the small size and/or spatial positioning of displaced chromosome fragments may pose a challenge for efficient reincorporation into daughter cell nuclei at the completion of mitosis. To visualize whether dispersed mitotic fragments accumulated within the interphase cytoplasm, we stained semi-permeabilized DLD-1 cells with a double-stranded DNA (dsDNA) antibody, which enabled a focus on the cytoplasm while minimizing intense nuclear DNA staining. Low levels of cytoplasmic dsDNA foci were triggered by micronucleation in WT cells, as we previously reported^39^. However, CIP2A KO cells exhibited elevated baseline levels of cytoplasmic dsDNA foci that were significantly exacerbated by micronucleus induction (**Fig. 4a**). To confirm that cytoplasmic dsDNAs originated from the Y chromosome, we performed interphase FISH on CIP2A KO cells; as expected, these dsDNAs were indeed specific for the Y chromosome but not a control X chromosome (**Fig. 4b-c**). Interestingly, most cytoplasmic dsDNAs (79% from sg3 and sg4 combined, *n* = 321 CIP2A KO cells) continued to harbor active γH2AX marks (**Fig. 4d**) persisting from chromosomal damage accrued within micronuclei during the previous cell cycle.

**Figure 4.**
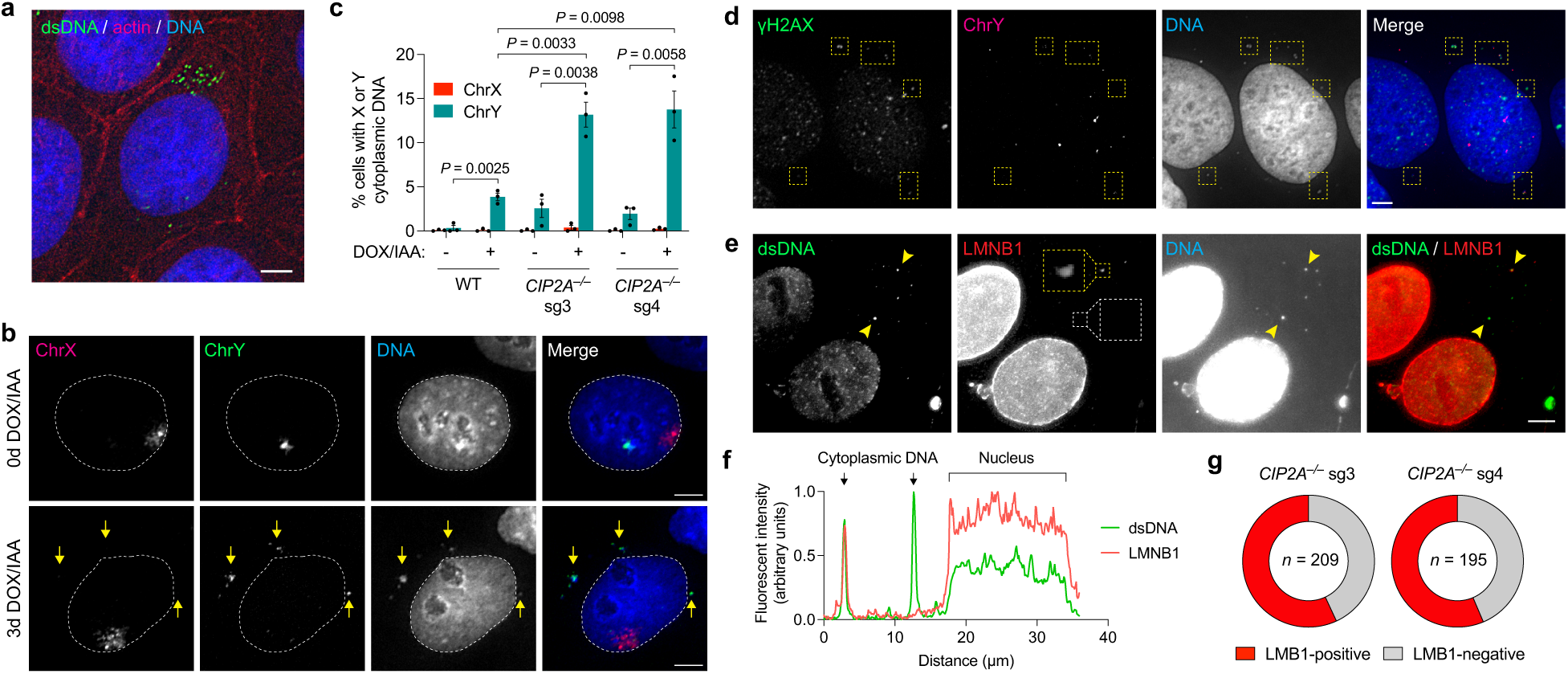
Dispersed chromosome fragments mis-accumulate in the cytoplasm as nuclear membrane-deficient cytoplasmic DNA. **a)** Semi-permeabilized CIP2A sg3 KO DLD-1 cells treated with DOX/IAA and stained with a double-stranded DNA (dsDNA) antibody. Actin staining is used to visualize the perimeter of the cell membrane. Scale bar, 5 μm. **b)** DNA FISH using X and Y chromosome paint probes showing Y chromosome-specific cytoplasmic DNA foci in CIP2A sg3 KO cells following DOX/IAA treatment. Scale bar, 5 μm. **c)** Quantification of X and Y chromosome-specific cytoplasmic DNA foci with and without DOX/IAA induction from (**b**). Data represent mean ± SEM; indicated *P*-values were obtained by two-tailed t-test; *n* = 3 independent experiments; WT (-DOX/IAA: *n* = 2,410, +DOX/IAA: *n* = 3,322), CIP2A KO sg3 (-DOX/IAA = 1,960, +DOX/IAA = 2,946), CIP2A KO sg4 (-DOX/IAA = 2,100, +DOX/IAA = 3,940 cells). **d)** Examples of γH2AX-positive Y chromosome fragments (yellow boxes) in the cytoplasm of interphase CIP2A sg4 KO cells. Scale bar, 5 μm. **e)** Examples of cytoplasmic DNA foci that are positive (yellow box, see magnified inset) or negative (white box) for the nuclear membrane marker lamin B1. White arrow depicts line drawn for analyses in (**f**). Scale bar, 5 μm. **f)** Fluorescent intensity line scan analysis between the indicated arrows depicted in (**e**) showing examples of cytoplasmic DNA foci with and without co-localization of lamin B1. **g)** Proportion of cytoplasmic DNA foci with and without lamin B1 staining from (**f**). Pie charts represent mean; *n* = number of foci pooled from 2 independent experiments.

At mitotic exit, the nuclear envelope re-assembles around daughter cell genomes with the capacity to also re-form around individual chromosomes^40^. Small chromosome fragments positioned away from the genomic mass and/or mitotic spindle poles may encounter difficulties in establishing a proper nuclear membrane. To test this, we determined whether cytoplasmic dsDNAs were comprised of components of the nuclear membrane. Approximately half (57% and 56% in sg3 and sg4, respectively) of cytoplasmic DNA foci in CIP2A KO cells contained detectable lamin B1 at an abundance comparable to the nucleus (**Fig. 4e-f**). However, the remaining fraction of cytoplasmic dsDNAs were completely devoid of apparent lamin B1 (**Fig. 4e-g**), perhaps caused by defects in reassembling a nuclear membrane around short genomic fragments. Non-encapsulated cytoplasmic dsDNAs are potent activators of the cGAS-STING pathway to trigger an innate immune response^41^. Thus, in addition to serving as a source of cytoplasmic DNA following micronuclear envelope rupture^35,36,39^, our data show that micronuclei can generate a second wave of cytoplasmic DNA owing to failures in reincorporating displaced chromosome fragments into daughter cell nuclei.

### Balanced chromothripsis in cancer genomes

Fragment clustering biases the inheritance of shattered chromosome fragments from micronuclei toward a single daughter cell during mitosis (**Fig. 1c,d,g**). In cancer genomes, we predicted that a derivative chromosome generated by chromothripsis with minimal fragment loss would exhibit clusters of structural rearrangements lacking the DNA copy number oscillations characteristic of canonical chromothripsis^1,3,42,43^. Evidence of such chromothripsis events – termed balanced chromothripsis – has been reported in the germline^44,45^ and lung and prostate adenocarcinomas^46,47^. However, the patterns, frequencies, and consequences of balanced chromothripsis in cancer genomes remain largely unknown due to the commonly used requirement of detecting DNA copy number oscillations to call chromothripsis events^3,42^.

To explore this concept, we used ShatterSeek (Cortes-Ciriano et al., 2020) to analyze whole-genome sequencing data from 2,575 tumors spanning 37 cancer types from the Pan-Cancer Analyses of Whole Genomes (PCAWG) Consortium for evidence of balanced chromothripsis (see **Methods**). Applying a strict threshold for zero to minimal DNA copy number changes, high-confidence balanced chromothripsis events were detected in ∼5% of the cancer genomes analyzed (119 of 2,575 tumor samples) (**Fig. 5a**). In the PCAWG cohort, the highest frequency of balanced chromothripsis were found in prostate adenocarcinomas (19.6%), soft-tissue liposarcomas (15.8%) and bone osteosarcomas (11%). In prostate adenocarcinomas, these events were distinct from chromoplexy as inter-chromosomal translocations were absent from the clusters of rearrangements^3,48^. Balanced chromothripsis was found in more than one chromosome in 19 cases. Examples of canonical and balanced chromothripsis events are shown in **Fig. 5b-g** and **Extended Data Fig. 11**, which exhibit complex and localized rearrangements reminiscent of chromothripsis but without the characteristic oscillations in DNA copy number states. Among cancer genomes with a balanced chromothripsis event, 102 of 119 (∼86%) disrupted at least one gene and 23 (∼19%) harbored chromosome breakpoints in putative cancer driver genes, including the tumor suppressor *PTEN* (**Fig. 5f**), *FOXO3*, and *ARID2*. Additionally, balanced chromothripsis generated fusion genes in 10 samples that included established oncogenic fusions, such as *CCDC170–ESR1* in breast adenocarcinomas, *RAB3C–PDE4D* in both skin melanomas and prostate adenocarcinomas, *CCT5–FAM173B* in bladder transitional cell carcinoma, and *TMPRSS2-ERG* in prostate adenocarcinomas (**Fig. 5g**, see **Extended Data Table 1** for complete list). Together, these results show that balanced chromothripsis underpins the acquisition of cancer driver events across diverse tumor types.

**Figure 5.**
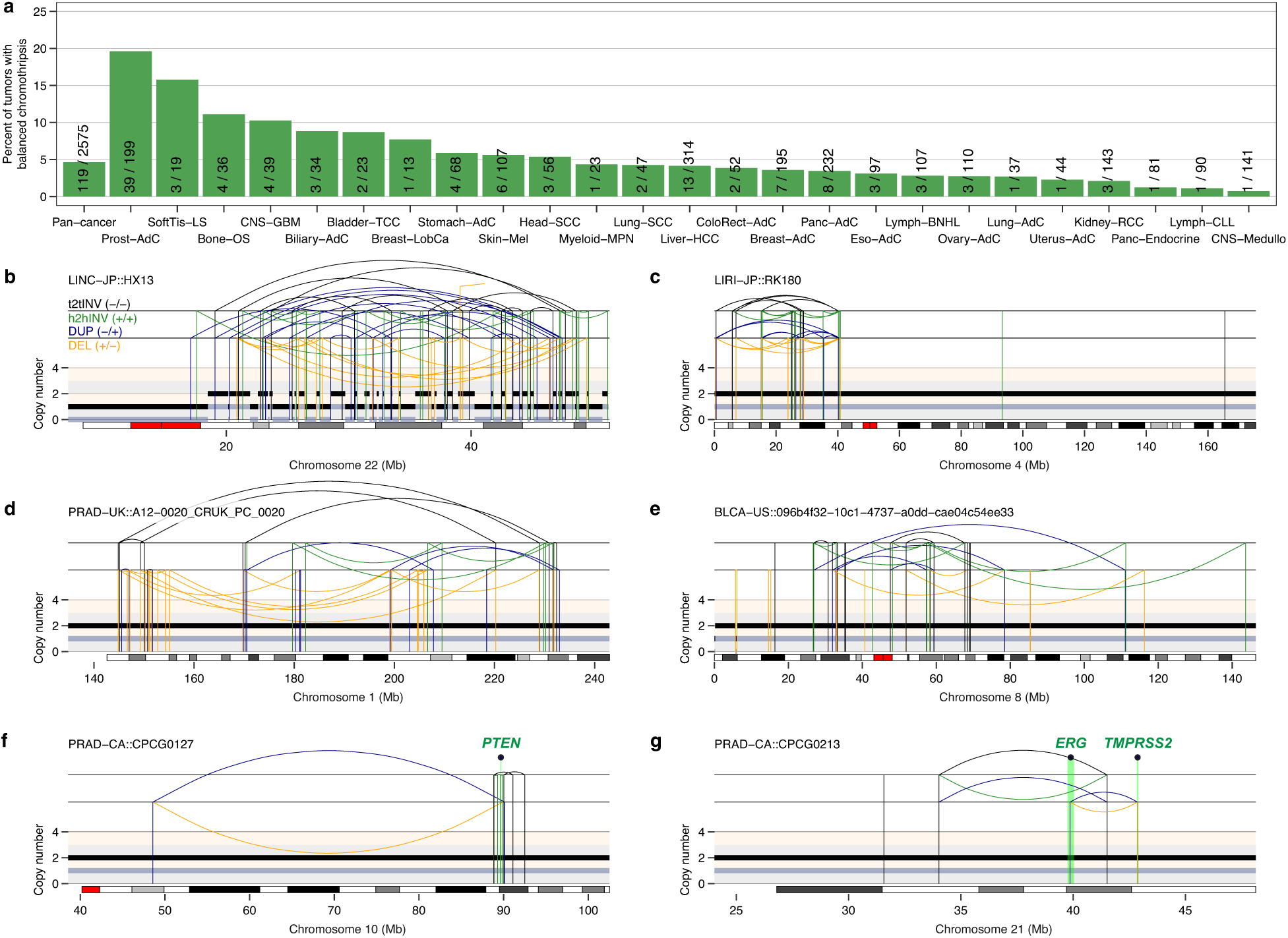
Prevalence of DNA copy number-neutral, balanced chromothripsis across diverse cancer types. **a)** Frequency of balanced chromothripsis in the ICGC/TCGA Pan-Cancer Analyses of Whole Genomes (PCAWG) cohort. Fractions represent the number of tumors with balanced chromothripsis in at least one chromosome over the total number of tumors of each type analyzed. **b)** Example of a canonical chromothripsis event affecting the q-arm of chromosome 22 in liver hepatocellular carcinoma with the characteristic pattern of DNA copy number oscillations. **c-e)** Examples of balanced chromothripsis events characterized by clusters of interleaved rearrangements, as expected for the random re-joining of genomic fragments shattered in chromothripsis, but without DNA loss, as indicated by the lack of deletions. **f-g)** Examples of balanced chromothripsis events causing inactivation of *PTEN* (**f**) and generating a *TMPRSS2-ERG* gene fusion (**g**) in two prostate adenocarcinomas. The cancer types shown in (**a**) are abbreviated as follows: Biliary-AdC, biliary adenocarcinoma; Bladder-TCC, bladder transitional cell carcinoma; Bone-Osteosarc, bone osteosarcoma; Breast-AdC, breast adenocarcinoma; CNS-GBM, central nervous system glioblastoma; CNS-Medullo, CNS medulloblastoma; ColoRect-AdC, colorectal adenocarcinoma; Eso-AdC, esophagus adenocarcinoma; Head-SCC, head-and-neck squamous cell carcinoma; Kidney-RCC, kidney renal cell carcinoma; Liver-HCC, liver hepatocellular carcinoma; Lung-AdC, lung adenocarcinoma; Lymph-BNHL, lymphoid mature B-cell lymphoma; Ovary-AdC, ovary adenocarcinoma; Panc-AdC, pancreatic adenocarcinoma; Panc-Endocrine, pancreatic neuroendocrine tumor; Prost-AdC, prostate adenocarcinoma; Skin-Mel, skin melanoma; Stomach-AdC, stomach adenocarcinoma. The total and minor copy-number data in (**c-f**) are represented in black and grey, respectively. DUP: duplication; DEL: deletion; t2tINV: tail-to-tail inversion; h2hINV: head-to-head inversion.

## Discussion

Here we identified a novel mechanism regulating the mitotic behavior of acentric DNA fragments arising from pulverized chromosomes in micronuclei. We propose a multi-step model (**Extended Data Fig. 12**) in which micronuclear envelope rupture initiates the diffusion and mis-localization of CIP2A into micronuclei during interphase, where it can prematurely associate with TOPBP1 to engage with DNA lesions. Upon nuclear envelope breakdown at mitotic entry, the CIP2A-TOPBP1 complex facilitates the clustering of chromosome fragments throughout mitosis. How CIP2A-TOPBP1 functions to tether fragments remain to be determined, but could occur through higher-order molecular interactions mediated by an extensive coiled-coil domain on CIP2A^30^ and/or the condensate-forming property of TOPBP1^49^. Mitotic clustering promotes the biased partitioning of most chromosome fragments *en masse* toward one daughter cell, which are then reincorporated into the interphase nucleus and persist as micronuclei bodies^28^ in the next cell cycle. Finally, clustered fragments that are spatially positioned in close nuclear proximity may become reassembled with increased DNA repair kinetics^50^ to generate a spectrum of genomic rearrangements^6^.

Mitotic clustering can safeguard against further genomic instability inflicted onto mis-segregated chromosomes; for example, by ensuring that most acentric fragments are inherited along with the centromere-containing fragment. Although this mechanism can minimize DNA copy number loss, we propose several non-mutually exclusive explanations for the loss of genomic fragments associated with chromothripsis. First, some fragments may fail to participate in clustering due to inefficiencies in tethering all acentric pieces. Additional factors may exist that promote or inhibit CIP2A-TOPBP1 activity, and it remains unclear whether this regulation differs across cell- and/or tissue-types. Second, micronuclear DNAs have been reported to exhibit under-replication during S-phase^5,12^, which can be caused by defective nucleocytoplasmic transport of the DNA replication machinery and/or the dilution of replication components following micronucleus rupture. Lastly, the loss of some fragments may arise from the inability of specific DNA repair pathways to re-assemble all chromosome fragments, which then become lost throughout subsequent rounds of cell division.

The biased inheritance of acentric fragments to a single daughter may explain the origins of balanced chromothripsis, which were detected in ∼5% of pan-cancer genomes. This is likely a conservative estimate as we applied a strict DNA copy number loss threshold to limit detection to high-confidence samples that may represent one extreme of a spectrum. The prevalence of canonical chromothripsis in cancer genomes likely reflects strong positive selection pressure owing to the increased risk of tumor suppressor deletions caused by partial chromosome loss^1^. In contrast, balanced chromothripsis – which relies on the precise location of rearrangement breakpoints to disrupt gene(s) – may be better tolerated in the germline and more capable of generating karyotypes that are compatible with organismal development^15^. These factors may in part contribute to the complex yet balanced rearrangement landscapes found in congenital disorders^16^. Further studies will be needed to define the specific cellular contexts in which these mechanisms are operative, as well as their contributions to cancer genome evolution and germline disorders.

## Materials and Methods

### Cell lines and reagents

DLD-1, HeLa, PC3 (a gift from Sihan Wu, UT Southwestern Medical Center, USA), HEK293T, and 293GP cells were cultured in DMEM (Gibco) containing 10% tetracyclin-free fetal bovine serum (Omega Scientific) and 100 U/ml penicillin-streptomycin. All cell lines were maintained at 37°C under 5% CO2 and atmospheric oxygen and were routinely confirmed free of mycoplasma contamination using the Universal Mycoplasma Detection Kit (ATCC).

Doxycycline (DOX) and auxin (indole-3-acetic acid, IAA) (Sigma-Aldrich) were dissolved in cell culture-grade water and used at 1 μg/mL and 500 μM, respectively. Geneticin (G418 Sulfate) and zeocin (InvivoGen) were used at selection concentrations of 300 and 50 μg/mL, respectively. For cell cycle arrest experiments, 100 ng/mL nocodazole (Sigma-Aldrich), 100 ng/mL colcemid (KaryoMAX, Thermo Fisher), or 10 μM MG132 (a gift from Joachim Seemann, UT Southwestern Medical Center, USA) were used for mitotic arrest, and 10 μM of the CDK1 inhibitor RO-3306 (Sigma) was used for G2 arrest, all of which were dissolved in dimethylsulfoxide (DMSO). 50 nM CENP-E inhibitor (GSK-923295) and 480 nM MPS1 inhibitor (NMS-P715, Cayman Chemical), were used to induce random chromosome mis-segregation and micronuclei formation in HeLa cells. For ionizing radiation experiments, cells were irradiated with γ-ray (2 Gy) generated by a Mark I ^137^Cs irradiator (JL Shepherd) and fixed one-hour post-irradiation for immunofluorescence analysis. Small interfering RNA (siRNA) transfections were conducted with Lipofectamine RNAiMAX reagent (Thermo Fisher). siRNAs were synthesized (Thermo Fisher) and used at a final concentration of 20 nM. All siRNAs used in this study are provided in **Supplementary Table 1**.

### Genome engineering of cell lines

To generate CIP2A knockout (KO) DLD-1 cells, target sequences for guide RNAs were designed using CRISPick (Broad Institute). Oligonucleotides encoding guide RNAs targeting exon 1 of CIP2A (denoted as sg3 and sg4, listed in **Supplementary Table 1**) were cloned into the BsmBI restriction site of the Lenti-Cas9-gRNA-TagBFP2 vector (Addgene 124774) and packaged in 293T cells by co-transfection with pMD2.G (Addgene 12259) and psPAX2 (Addgene 12260) using X-tremeGENE 9 (Sigma-Aldrich). Viral supernatants after 48- or 72-hour transfection were filtered (0.45 μm) and DLD-1 cells were infected in the presence of 5 μg/mL polybrene (Sigma-Aldrich) for ∼24 hours. Fluorescent cells were isolated by fluorescence-activated cell sorting (FACS) into 96-well plates (BD FACSAria II). KO clones were then expanded and verified by Sanger sequencing and immunoblotting.

To generate DLD-1 KO cells by ribonucleoprotein (RNP)-mediated CRISPR genome editing, two sgRNAs per gene were synthesized (Synthego) and co-transfected with TrueCut Cas9 protein v2 (Invitrogen) using Lipofectamine CRISPRMAX Cas9 transfection reagent (Invitrogen). All sgRNA sequences used in this study are provided in **Supplementary Table 1**. After transfection, cells were plated by limiting dilution into 96-well plates. Single cell-derived clones were expanded, screened by PCR for targeted deletions, and confirmed to harbor frameshift deletion mutations by Sanger sequencing. All PCR primers (Sigma) are provided in **Supplementary Table 1**.

For complementation experiments, CIP2A cDNA (a gift from Qing Zhang, UT Southwestern Medical Center, USA) and a HaloTag (Addgene 112852) were cloned into pBABE-zeo and packaged in 293GP cells by co-transfection with pVSV-G using X-tremeGENE 9. CIP2A KO cells generated by RNP-mediated gene editing were infected with retroviruses encoding full-length CIP2A or delta NES mutant (lacking amino acids 561-625) fused to an N-terminus HaloTag for 24 hours and selected with zeocin for 10 days. For expression of other exogenous genes, H2B-mCherry (a gift from Hongtao Yu, UT Southwestern Medical Center, USA) and cGAS-GFP constructs (a gift from Zhijian Chen, UT Southwestern Medical Center, USA) were used to generate viruses for transduction of DLD-1 cells, as described.

### Chromosome labeling using dCas9-SunTag

To label the Y chromosome in live cells, the SunTag labeling system was adopted and modified as described below. DYZ1 repeats (3,584 bp, sequence information provided by Helen Skaletsky and David Page, Whitehead Institute, HHMI, USA) was analyzed by CRISPick (Broad Institute) and five sgRNA sequences were selected for targeting DYZ1 repeats. scFv-GCN4-sfGFP-GB1-NLS from SunTag plasmid (Addgene 60906) was cloned into a lentiGuide-puro vector (Addgene 52963). Lentiviral supernatants, which were packaged in 293T cells by co-transfection with pMD2.G and psPAX2 with either lentiGuide-scFv-GCN4-sfGFP-GB1-NLS or pHRdSV40-dCas9-10xGCN4_v4-P2A-BFP (SunTag plasmid Addgene 60903), and retroviral supernatants, which were packaged in 293GP cells by co-transfection of pBABE-H2B-mCherry with pVSV-G after 48- or 72-hour transfection, were filtered (0.45 μm) and DLD-1 cells were infected in the presence of 5μg/mL polybrene (Sigma-Aldrich) for ∼24 hours. Fluorescent cells were isolated by FACS (BD FACSAria II) and plated by limiting dilution into 96-well plates. Single cell-derived clones were expanded and screened for expected SunTag signals.

### Live-cell imaging

DLD-1 cells expressing dCas9-SunTag and H2B-mCherry were plated in Nunc Lab-Tek chambered coverglasses. Images were acquired on a DeltaVision Ultra microscope (GE Healthcare) in a humidity- and temperature-controlled (37°C) environment supplied with 5% CO2 at 5 min. intervals for 16 hours using a 60x objective with 11 × 0.5 μm z-sections under low power exposure. For CIP2A-HaloTag imaging, cells were labeled with 200 nM JF646 ligand (Promega) for 15 min. and washed with fresh media before imaging. Images were deconvolved and maximum intensity quick projections were generated using softWoRx. Videos were analyzed using Fiji.

### Immunofluorescence

DLD-1 cells were plated onto CultureWell gaskets (Grace Bio-Labs) and assembled glass slides were fixed with 4% formaldehyde for 10 min. For dispersion analysis, cells were arrested in mitosis for 4 hours using colcemid and collected by shake-off. Cell suspensions were concentrated to 1 × 10^6^ cells/mL in PBS and centrifuged onto glass slides using a Cytospin 4 cytocentrifuge (Thermo Scientific). Fixed cells were permeabilized with 0.3% Triton X-100 in PBS for 5 min and incubated with Triton Block (0.2 M glycine, 2.5% fetal bovine serum, 0.1% Triton X-100, PBS). The following primary antibodies were used at the indicated dilutions in Triton Block: 1:500 anti-CIP2A (Santa Cruz, sc-80659), 1:500 anti-TOPBP1 (Santa Cruz, sc-271043), 1:300 anti-TOPBP1 (Millipore, ABE1463), 1:1000 anti-phospho H2AX (S139) (Millipore, 05-636), 1:1000 anti-phospho H2AX (S139) (Cell Signaling, 2577) antibodies. Cells were washed with 0.1% Triton X-100 in PBS, incubated with 1:1000 dilutions of Alexa Fluor-conjugated secondary antibodies (Invitrogen), and washed with 0.1% Triton X-100. Immunofluorescence-labeled cells were fixed with Carnoy’s fixative for 15 min and rinsed with 80% ethanol. Air-dried cells were then used for DNA fluorescence in situ hybridization (FISH), as described below.

For micronuclei analysis, DLD-1 and HeLa cells were grown on glass coverslips and fixed with PTEMF (0.2% Triton X-100, 0.02 M PIPES pH 6.8, 0.01 M EGTA, 1 mM MgCl2, and 4% formaldehyde) for 10 mins., followed by two washes in 1X PBS. Samples were blocked with 3% bovine serum albumin diluted in PBS. Cells were incubated for 1 hour at room temperature with the following primary antibodies diluted in 3% BSA: 1:500 anti-CIP2A (sc-80659, Santa Cruz), 1:500 anti-TOPBP1 (sc-271043, Santa Cruz), 1:1000 anti-phospho-histone H2AX (Ser139, 2577, Cell Signaling), and 1:1000 anti-cGAS (15102, Cell Signaling). After 3 × 5 min. washes, fluorescence-conjugated secondary antibodies (Thermo Fisher Scientific) were diluted 1:1000 in 3% BSA and applied to cells for 1 h at room temperature, followed by 2 × 5 min. washes with 1X PBS. DNA was counterstained with DAPI and cells were mounted in ProLong Gold antifade mounting solution.

For cytosolic dsDNA staining, cells were fixed with 4% formaldehyde for 10 min and then treated with 0.02% saponin in PBS for 5 min. Semi-permeabilized cells were incubated with blocking solution (2.5% fetal bovine serum in PBS) and followed by incubation with anti-dsDNA antibody (1:250 in blocking solution, sc-58749, Santa Cruz) at 4°C overnight. After washing with PBS, cells were incubated with 1:1000 dilutions of Alexa Fluor-conjugated secondary antibodies (Invitrogen) in blocking solution for 1 h and washed with PBS. Cells were then fully permeabilized with 0.3% Trion X-100 in PBS for 5 min and washed with PBS. Permeabilized cells were incubated with 5 U/mL of fluorescent phalloidin (Biotium) in PBS for 20 min and washed with PBS.

### Metaphase spread preparation

Cells were treated with 100 ng/mL colcemid (KaryoMAX, Thermo Fisher) for 4-5 hours before harvesting by trypsinization and centrifugation. Cell pellets were resuspended in 500 μL PBS followed by adding 5 mL of 75 mM KCl solution dropwise while gently vortexing. Cells were incubated for 6 min in 37°C water bath and fixed using freshly prepared Carnoy’s fixative (3:1 methanol:acetic acid), followed by centrifugation and resuspension in Carnoy’s fixative. Cells were subsequently dropped onto slides and air dried.

### DNA fluorescence in situ hybridization (FISH)

DNA FISH probes (MetaSystems) were applied to metaphase spreads and sealed with a coverslip using rubber cement. Slides were co-denatured on a heat block at 75°C for 2 min. and then hybridized at 37°C in a humidified chamber overnight. The next day, coverslips were removed, and the slides were washed with 0.4X SSC at 72°C for 2 min. and rinsed with 2X SSC with 0.05% Tween-20 at room temperature for 30 seconds. After washing, slides were counterstained with DAPI, air dried, and mounted in ProLong Gold antifade mounting solution.

### Fixed-cell microscopy

Immunofluorescence images were captured on a DeltaVision Ultra (GE Healthcare) microscope system equipped with 4.2 MPx sCMOS detector. Interphase nuclei and micronuclei images were acquired with a 100x objective (UPlanSApo, 1.4 NA) and 1 × 0.2-μm z-section. Quantitative fluorescence image analyses were performed using Fiji. IF-FISH images were acquired with a 60x objective (PlanApo N 1.42 oil) and 15 × 0.2 μm z-sections. Deconvolved maximum intensity projections were generated using softWoRx program.

Metaphase FISH images were acquired on a Metafer Scanning and Imaging Platform microscope (MetaSystems). Slides were first scanned for metaphases using M-search with a 10x objective (ZEISS Plan-Apochromat 10x/0.45), and metaphases were automatically imaged using Auto-cap with a 63x objective (ZEISS Plan-Apochromat 63x/1.40 oil). Images were analyzed using Isis Fluorescence Imaging Platform (MetaSystems).

### Chromosome inheritance between daughter pairs

Cells were seeded in 4-well chamber slides and treated with or without DOX/IAA for 48 hours. Cells were then arrested in G2 with 10 µM CDK1 inhibitor RO-3306 (Sigma) for 16 hours and released by washing with PBS for 3 times and adding fresh medium. After 90 mins, cells were fixed with 4% formaldehyde followed by IF FISH, as described. X and Y chromosome paint probes (MetaSystems) were used for FISH. For analysis of chromosome inheritance between daughter cells, pairs of newly formed daughter cells were imaged on a DeltaVision Ultra (GE Healthcare) microscope system. Images were split into separate channels for quantification using the ImageJ plugin Segmentation (Robust Automatic Threshold Selection) to create a mask for the FISH signals. Particles of the mask were analyzed to generate a list of regions of interest (ROI) for intensity measurements. FISH signal intensities were then measured in each pair of daughter cells for both the X and Y chromosomes. The distribution of FISH signal was calculated by the ratio of the daughter cell with the lower signal compared to the daughter cell with the higher signal.

### Chromosome fragment dispersion

Metaphase spreads were prepared as described and hybridized to Y chromosome paint probes (MetaSystems). Metaphases with fragmented Y chromosomes were identified and split into separate channels. Fragment dispersion was analyzed using the ImageJ plugin HullAndCircle to measure the convex hull of the Y chromosome fragments relative to all DAPI-stained chromosomes. Dispersion indices were calculated by dividing the area of Y chromosome fragments by the overall DAPI area followed by min-max normalization of all data points within each sample.

### Cell cycle profile

Cells were trypsinized, washed with PBS, and fixed with 70% ethanol in PBS at -20°C for 2 hours. Fixed cells were washed with PBS twice and incubated with staining solution (0.1 mg/mL RNase A, 0.1% Triton X-100, 10 μg/mL propidium iodide). Cells were analyzed using a FACSCalibur (BD Biosciences) flow cytometer.

### Immunoblotting

Whole-cell extracts were collected in SDS sample buffer and boiled for

5 min. Samples were resolved by SDS polyacrylamide gel electrophoresis, transferred to polyvinylidene fluoride membranes, and blocked with 5% milk diluted in PBST (PBS, 0.1% Tween-20). The following primary antibodies were diluted in PBST and used: 1:1000 anti-CIP2A (Santa Cruz, sc-80659), 1:5000 anti-α-tubulin (Cell Signaling, 3873), anti-TOPBP1 (Santa Cruz, sc-271043). Blots were incubated with 1:4000 dilutions of HRP-conjugated secondary antibodies (Invitrogen), incubated with SuperSignal West Pico Plus chemiluminescent substrate (Thermo Scientific), and processed using a ChemiDoc MP imaging system (Bio-Rad).

### Whole-genome sequencing analyses

To detect copy-number balanced chromothripsis events, we applied ShatterSeek v1.1^3^ (https://github.com/parklab/ShatterSeek) to 2,575 tumor-normal pairs from PCAWG that passed quality control criteria. We considered all chromosomes with a cluster of at least 5 structural variants (SVs). We considered all clusters irrespective of the number of copy number oscillations in the cluster. To call a cluster of SVs a copy-number-balanced chromothripsis event, we required: (*i*) at least 5 intra-chromosomal SVs; (*ii*) no translocation mapping to the genomic region encompassed by the cluster of SVs; we included this filter to distinguish balanced chromothripsis from chromoplexy events, which are characterized by chains of inter-chromosomal SVs with limited genomic DNA loss and thus could be misclassified as balanced chromothripsis if this filter was not applied; (*iii*) no overlap with chromoplexy calls generated for these tumors using ChainFinder^48^ as previously reported^3^; and (*iv*) that less than 1% of the genomic region encompassed by the cluster of SVs shows a copy number below the modal copy number of the chromosome. We applied this filter to ensure that balanced chromothripsis calls do not contain canonical chromothripsis events. Finally, all cases that passed these filters were examined manually by visualizing genomic rearrangement plots.

To find gene disruptions within the balanced chromothripsis clusters, we first downloaded gene coordinates from Ensembl^51^ (GRCh37) using biomaRt^52^. Next, we intersected the coordinates of the breakpoints and genes using bedtools^53^. We considered a gene to be disrupted if a breakpoint mapped within the region defined by the start and end coordinates of the gene ± 5 kilobases. We determined putative cancer driver genes using the pan-cancer driver catalog from the Hartwig Medical Foundation whole-genome sequencing cancer analysis pipeline (https://github.com/hartwigmedical/hmftools/blob/master/purple/DriverCatalog.md).

### Statistical analyses

Statistical tests were performed as described in the figure legends using GraphPad Prism version 9.1.0. Sample sizes, statistical analyses, significance values are described in the figure legends, denoted in the figure panel, or reported in the text. *P* values ≤ 0.05 were considered to be statistically significant. Error bars represent standard error mean unless otherwise stated.

## Author Contributions and Notes

Y.F.L and P.L conceived the project. Y.F.L., A.M., Q.H., and P.L. designed the experiments. Y.F.L, A.M., Q.H., E.G.M., A.G., J.T.S., J.E., G.N., and D.B. conducted cell biological experiments and analyzed the data. J.E.V.-I. and I.C.-C. performed analysis of whole-genome sequencing data. S.F.B. provided critical input. P.L. wrote the manuscript with input from all the authors.

This article contains supporting information online (Extended Data Figures 1-12, Extended Data Table 1, and Supplementary Table 1).

## Acknowledgements

We thank Helen Skaletsky and David Page for advice on targeting the DYZ1 array; Khuloud Jaqaman for advice on imaging analysis; Kevin Dean for assistance with microscopy; Zhijian Chen, Joachim Seemann, Sihan Wu, Hongtao Yu, and Qing Zhang for sharing reagents; and members of the Ly laboratory for insightful discussions. This work was supported by the Cancer Prevention and Research Institute of Texas (RR180050 to P.L.; RP21004 to E.G.M.), U.S. National Institutes of Health (R00CA218871 to P.L.; T32CA124334 to J.T.S.), and American Cancer Society Institutional Research Grant (ACS-IRG-21-142-16 to P.L.). I.C.-C. and J.E.V.-I. acknowledge the European Molecular Biology Laboratory for funding.

