## Extended Data Figures 1-12 for "Mitotic clustering of pulverized chromosomes from micronuclei"

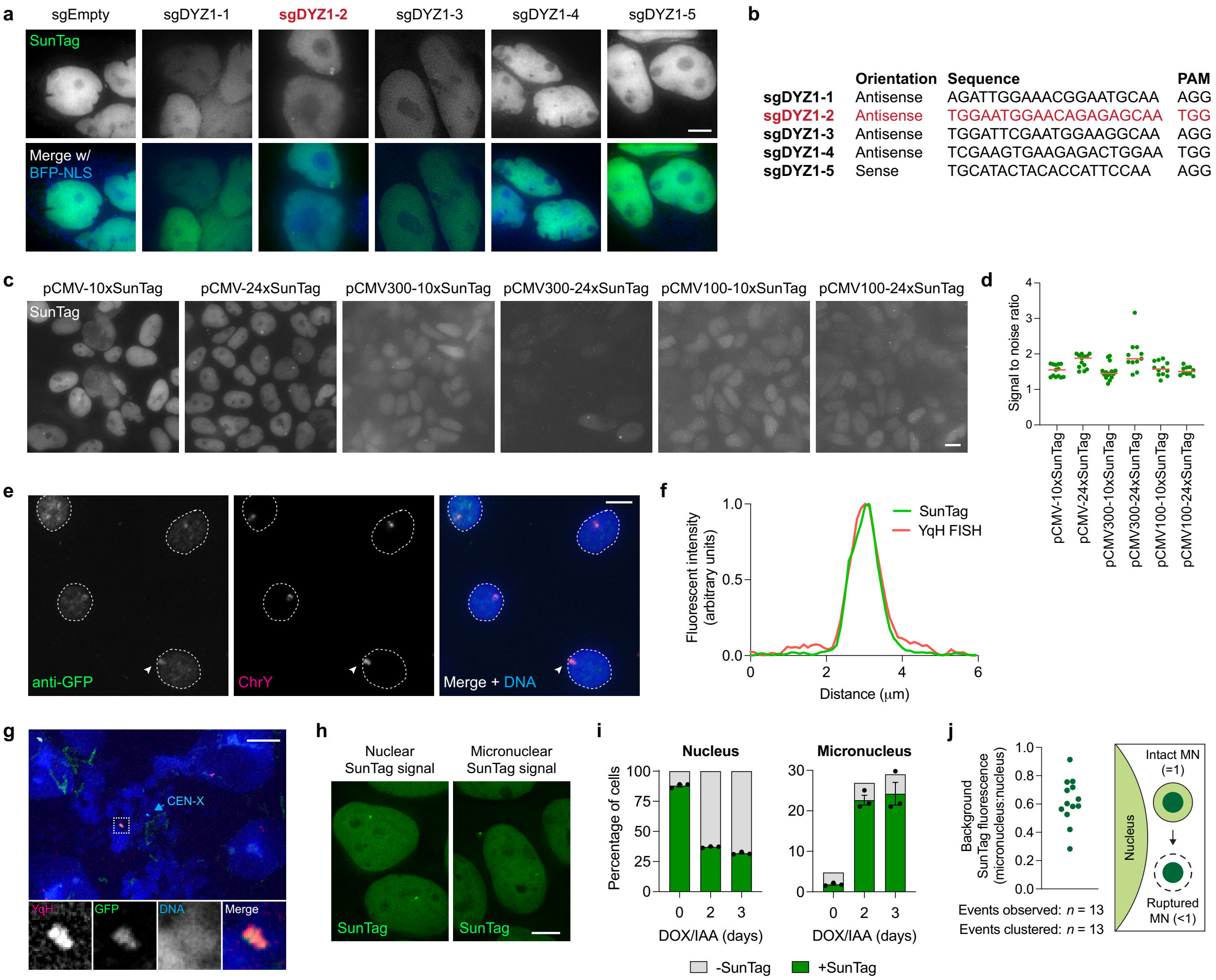


**Extended Data Figure 1. Development of a live-cell Y chromosome-labeling system by targeting dCas9-SunTag to the DYZ1 array. a)** Images of DLD-1 cell populations expressing dCas9-SunTag and sfGFP-scFv with the indicated sgRNAs targeting the DYZ1 array. Scale bar, 5 μm. **b)** List of sgRNA sequences used in (**a**). sgDYZ1-2 was used for the remainder of the study. **c)** Images of DLD-1 cell populations expressing sfGFP-scFv under the control of full-length or truncated CMV promoters with dCas9-SunTag containing the indicated scaffold lengths. Scale bar, 5 μm. **d)** Signal-to-noise measurements for the conditions shown in (**c**). Data represent mean of the indicated number of cells. **e)** IF-FISH image of interphase cells showing co-localization between an anti-GFP antibody recognizing sfGFP bound to dCas9-SunTag and DNA FISH probes targeting the Y chromosome q-arm heterochromatic array (YqH). Scale bar, 5 μm. **f)** Fluorescent line scan analysis of the indicated region marked in (**e**) showing high specificity of the SunTag with YqH. **g)** IF-FISH image of mitotic chromosomes showing co-localization between an anti-GFP antibody recognizing dCas9-SunTag with chromosome paint probes targeting YqH. Scale bar, 10 μm. **h)** Example images of live DLD-1 cells with dCas9-SunTag signals in the nucleus or micronucleus. Scale bar, 5 μm. **i)** Proportion of nuclei and micronuclei with or without dCas9-SunTag signals following DOX/IAA induction for the indicated number of days. Data represent mean ± SEM of $n$ = 3 independent experiments; 0 days = 1,044, 2 days = 1,070, 3 days = 1,123 cells. **j)** Background fluorescence measurements of non-dCas9-SunTag-bound sfGFP-scFv from $n$ = 13 micronuclei obtained from independent experiments (left) and schematic of intact and ruptured micronuclei measurements (right).

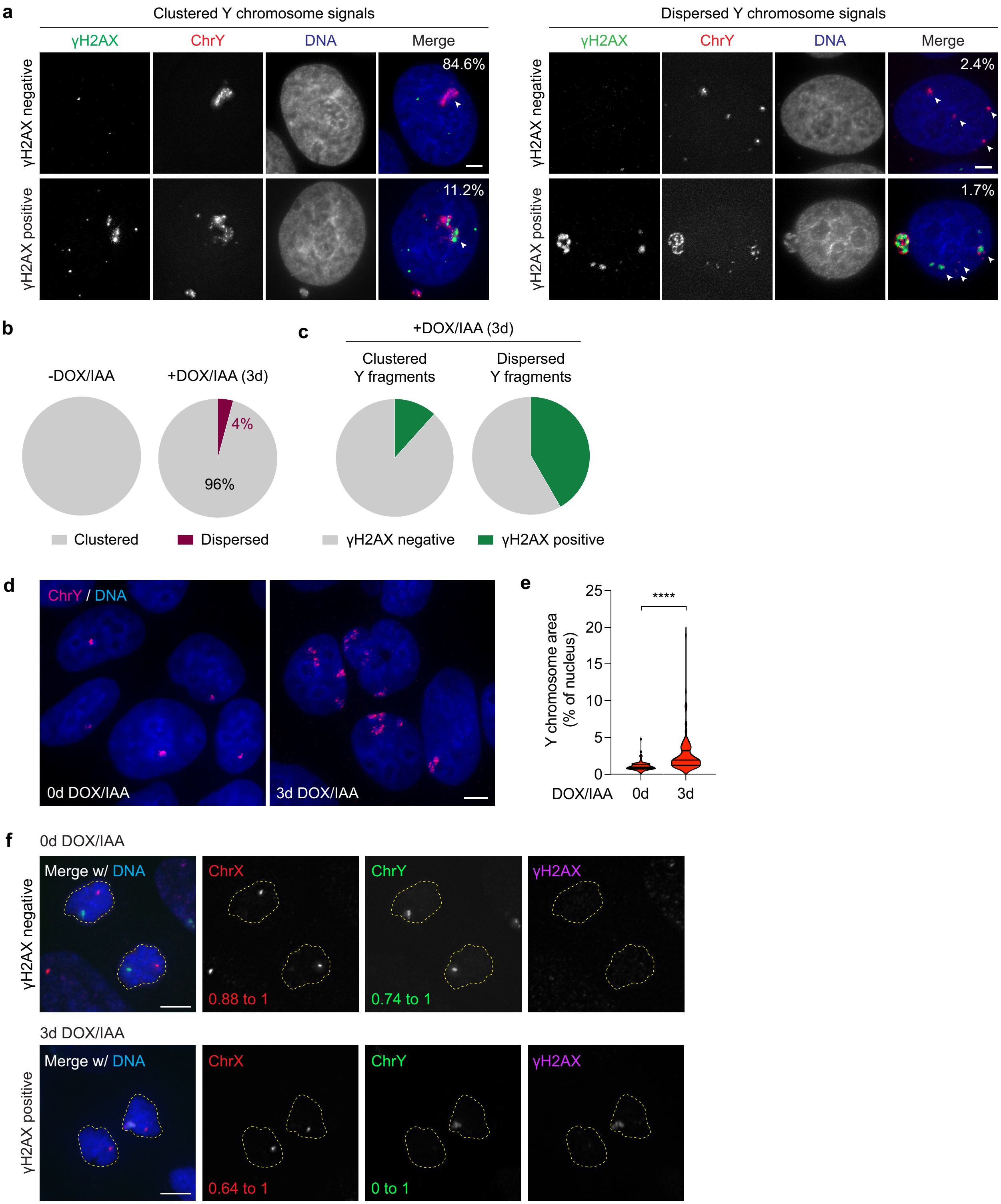


**Extended Data Figure 2. Micronucleation triggers clusters of damaged chromosome fragments that unevenly reincorporates into daughter cell nuclei. a)** Examples of clustered (top panels) and dispersed (bottom panels) Y chromosome signals in the interphase nucleus with or without γH2AX. Percentages represent the proportion of cells that exhibit each category following 3d DOX/IAA treatment. Scale bar, 5 μm. **b)** Pie charts depicting the fraction of control or DOX/IAA-treated cells with clustered or dispersed Y chromosome fragments during interphase. Data represent mean; -DOX/IAA: *n* = 236, +DOX/IAA: *n* = 573 cells. **c)** Pie charts depicting the γH2AX status of clustered and dispersed Y chromosome fragments. Data represent mean; clustered: *n* = 549, dispersed: *n* = 24 fragments. **d)** Re-integrated Y chromosome fragments occupy a larger nuclear space increased nuclear fluorescence signal area following DOX/IAA treatment. Scale bar, 5 μm. **e)** Violin plot quantification of (**d**) measuring Y chromosome FISH area over the total area of the nucleus, as indicated by DAPI staining; data pooled from 0d: *n* = 103, 3d: *n* = 105 Y chromosome clusters; ****$P \leq 0.0001$ by Welch's unpaired t-test. **f)** Images of equal segregation of an intact Y chromosome (top) and pulverized Y chromosome exhibiting unequal partitioning between daughter cells. Scale bar, 5 μm. Quantification shown in **Fig. 1g**.

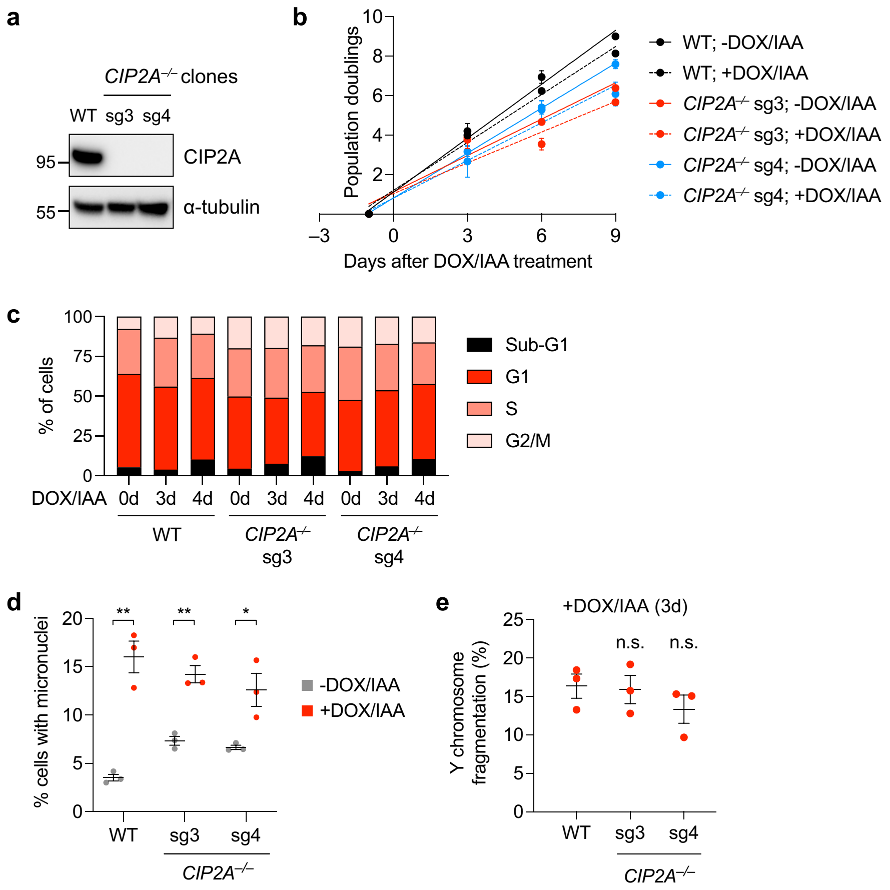


**Extended Data Figure 3. Characterization of CIP2A knockout DLD-1 cells. a)** Western blot confirmation of CIP2A KO clones. **b)** Growth curves of WT and CIP2A KO cells with and without DOX/IAA treatment over the indicated number of days. Data represent mean ± SEM; *n* = 3 replicates. **c)** Flow cytometry analysis of propidium iodide-stained WT and CIP2A KO cells showing similar cell cycle distribution profiles. **d)** Proportion of micronucleated cells with or without 2d DOX/IAA induction, as determined by DAPI staining. Data represent mean ± SEM; WT: ***P* = 0.0017, sg3: ***P* = 0.0023, sg4: **P* = 0.0261 by two-tailed t-test compared to untreated controls; *n* = 3 independent experiments; WT (-DOX/IAA = 5,521, +DOX/IAA = 3,718), CIP2A KO sg3 (-DOX/IAA = 3,436, +DOX/IAA = 2,450), CIP2A KO sg4 (-DOX/IAA = 3,999, +DOX/IAA = 2,930 cells). **e)** Frequency of Y chromosome fragmentation among Y chromosome-positive metaphase spreads following 3d DOX/IAA induction. Data represent mean ± SEM; not significant (ns), *P* > 0.05 by two-tailed t-test compared to WT controls; *n* = 3 independent experiments; WT = 234, CIP2A KO sg3 = 269, CIP2A KO sg4 = 284 cells.

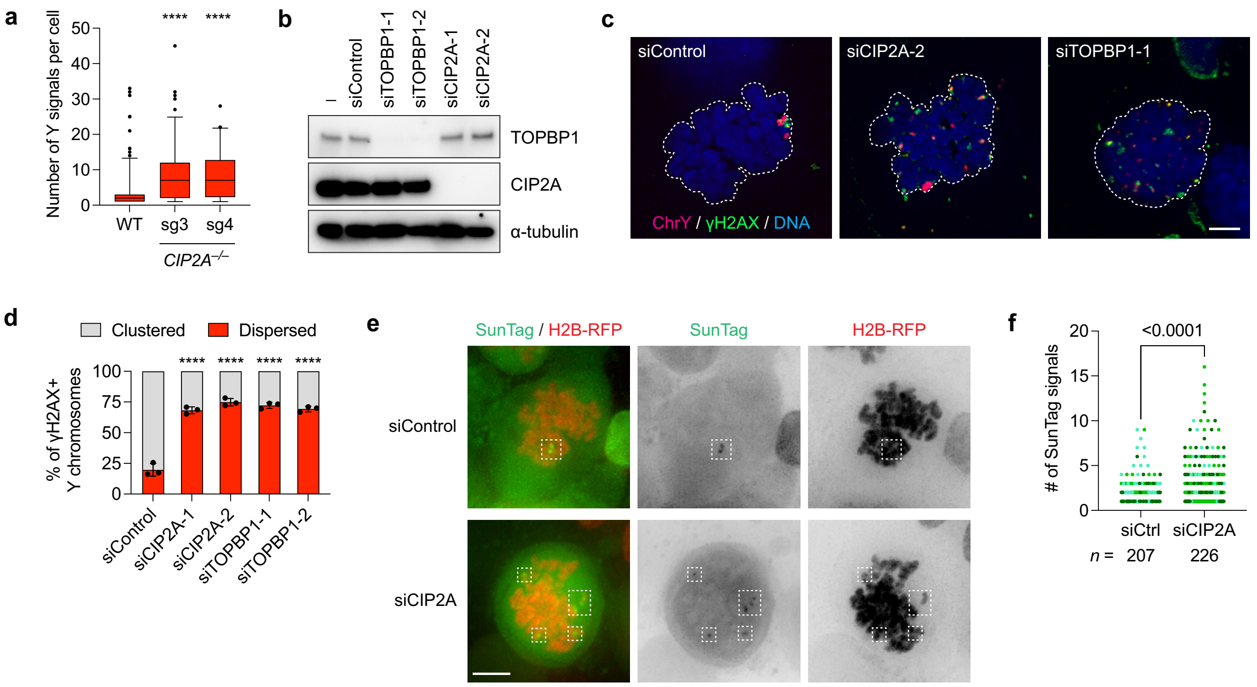


**Extended Data Figure 4. Loss of CIP2A-TOPBP1 disperses fragmented chromosomes in intact cells. a)** Quantification of Y chromosome-positive signals in WT and CIP2A KO cells. Data represent median with 5-95 percentiles; ****$P \leq 0.0001$ by two-tailed t-test compared to WT controls; WT: $n$ = 214, CIP2A KO sg3: $n$ = 113, CIP2A KO sg4: $n$ = 84 cells pooled from 2 independent experiments. **b)** Western blot of CIP2A and TOPBP1 depletion in WT DLD-1 cells using two independent small interfering RNAs. **c)** Images of fragmented Y chromosomes in mitotic WT DLD-1 cells depleted of CIP2A or TOPBP1 prior to DOX/IAA induction. Scale bar, 5 μm. **d)** Quantification of fragment clustering and dispersion from (**b**). Data represent the mean ± SEM of $n$ = 3 independent experiments; ****$P \leq 0.0001$ by one-way ANOVA with multiple comparisons test compared to siControl sample; siControl = 128, siCIP2A-1 = 145, siCIP2A-2 = 178, siTOPBP1-1 = 138, siTOPBP1-2 = 133 cells. **e)** Live-cell images of dCas9-SunTag signals from nocodazole-arrested DLD-1 cells showing increased SunTag-positive fragments following CIP2A depletion. Scale bar, 5 μm. **f)** Quantification of the number of SunTag-positive signals from (**d**). Individual data points represent a single cell; data pooled from 3 independent experiments; unpaired t-test with Welch's correction.

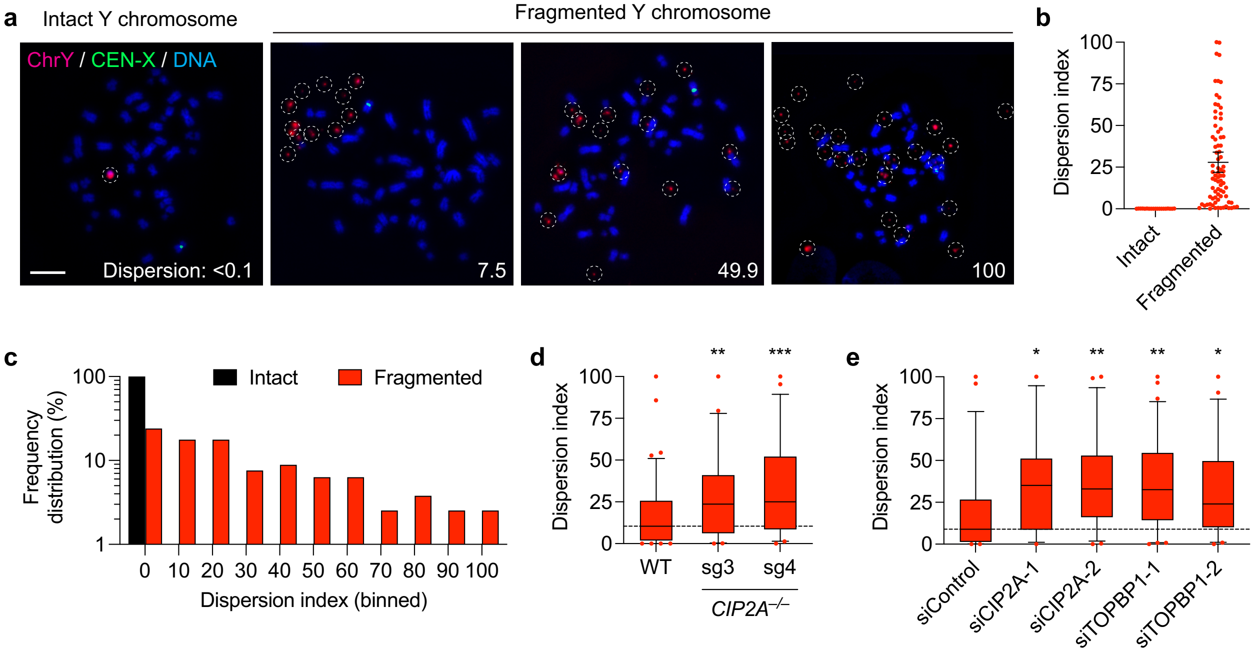


**Extended Data Figure 5. Loss of CIP2A-TOPBP1 disperses fragmented chromosomes on metaphase spreads. a)** Metaphase spreads were collected from DLD-1 cells treated with DOX/IAA and hybridized to Y chromosome FISH probes. Examples of intact and fragmented Y chromosomes are shown along with dispersion index (see **Methods** for measurements). Scale bar, 5 $\mu$m. **b)** Quantification of metaphase fragment dispersion from (**a**). Data represent individual metaphase spreads with an intact or fragmented Y chromosome; intact: $n$ = 19, fragmented: $n$ = 79 metaphases from 3 independent experiments. **c)** Distribution of dispersion indices for intact and fragmented Y chromosomes from data shown in (**b**); intact: $n$ = 19, fragmented: $n$ = 79 metaphases from 3 independent experiments. **d)** CIP2A KO cells exhibit increased fragment dispersion on metaphase chromosome spreads. Data represent median with 5-95 percentiles; $^{**}P$ = 0.0042, $^{***}P$ = 0.0002 by one-way ANOVA with multiple comparisons test compared to WT control sample; WT: $n$ = 83, sg3: $n$ = 54, sg4: $n$ = 44 metaphases from 3 independent experiments. **e)** CIP2A or TOPBP1 depletion increases fragment dispersion on metaphase chromosome spreads. Data represent median with 5-95 percentiles; $^{*}P \leq 0.05$; $^{**}P \leq 0.01$ by one-way ANOVA with multiple comparisons test compared to siControl sample; siControl: $n$ = 57, siCIP2A-1: $n$ = 38, siCIP2A-2: $n$ = 52, siTOPBP1-1: $n$ = 63, siTOPBP1-2: $n$ = 49 metaphases from 3 independent experiments.

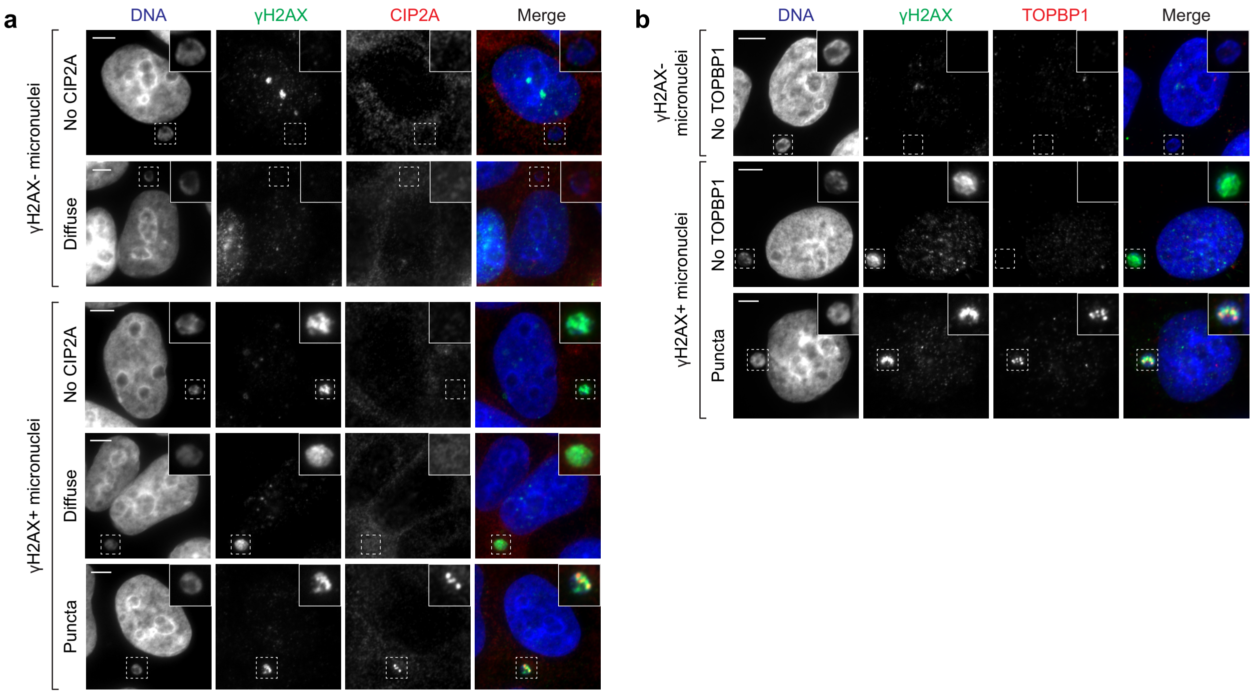


**Extended Data Figure 6. Interphase recruitment of CIP2A-TOPBP1 to γH2AX-positive ruptured micronuclei in DLD-1 cells. a-b)** Examples of distinct localization patterns of CIP2A (**a**) or TOPBP1 (**b**) in intact (γH2AX-negative) and ruptured (γH2AX-positive) micronuclei. Quantification shown in **Fig. 3a-b**. Scale bar, 10 μm.

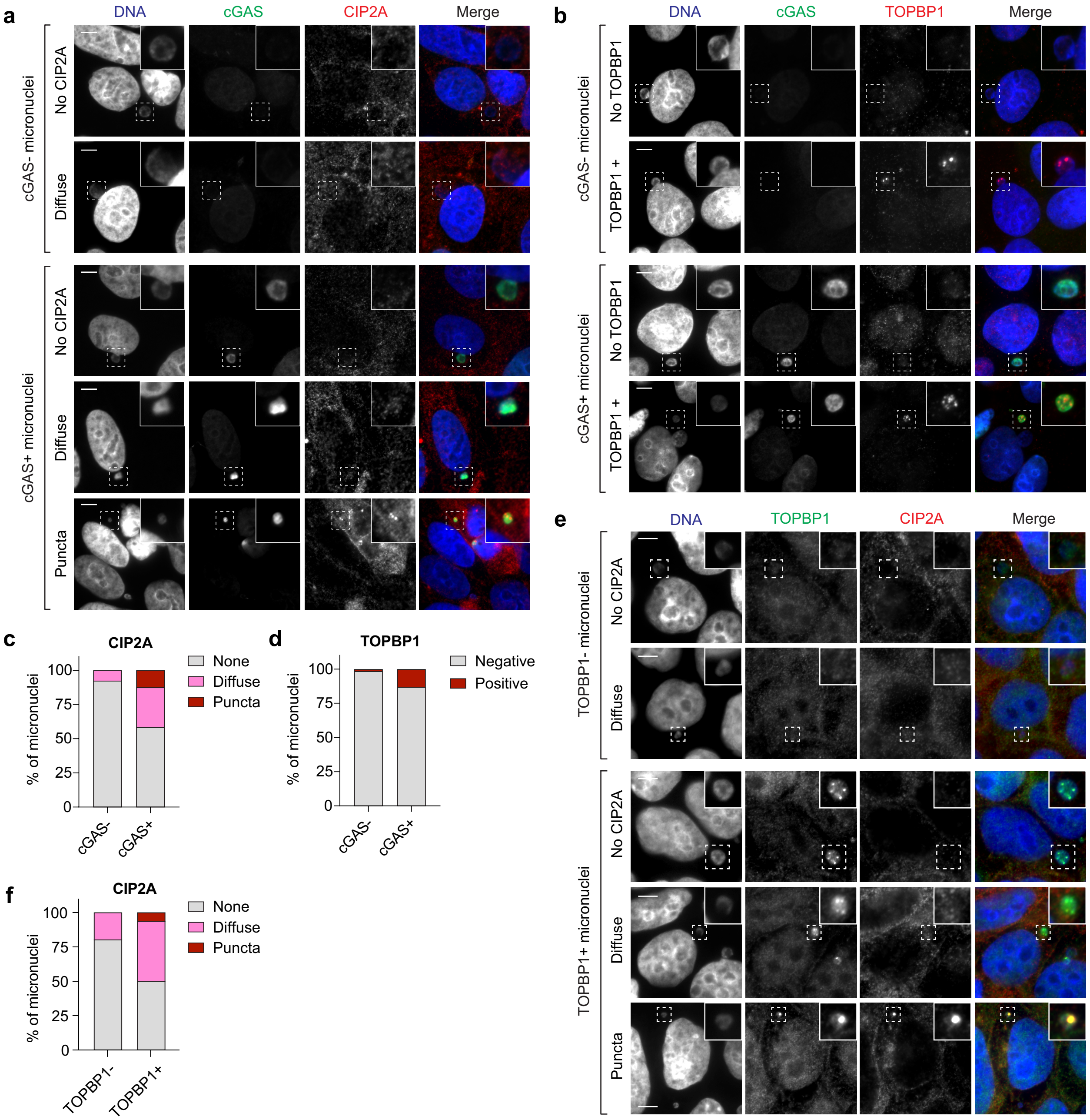


**Extended Data Figure 7. Interphase CIP2A-TOPBP1 patterns in cGAS-positive ruptured micronuclei in DLD-1 cells. a-b)** Examples of distinct localization patterns of CIP2A (**a**) or TOPBP1 (**b**) in intact (cGAS-negative) and ruptured (cGAS-positive) micronuclei from DLD-1 cells expressing exogenous cGAS-GFP. Scale bar, 10 μm. **c)** Frequency of CIP2A localization patterns in intact (cGAS-negative) and ruptured (cGAS- positive) micronuclei from (**a**). Data represent mean; cGAS-negative: *n* = 52, cGAS-positive: *n* = 48 micronuclei. **d)** Frequency of TOPBP1 localization patterns in intact (cGAS-negative) and ruptured (cGAS-positive) micronuclei from (**b**). Data represent mean; cGAS-negative: *n* = 62, cGAS-positive: *n* = 38 micronuclei. **e)** Localization patterns of CIP2A and TOPBP1 in DOX/IAA-treated DLD-1 cells. Scale bar, 10 μm. **f)** Frequency of CIP2A localization patterns in TOPBP1-negative and TOPBP1-positive micronuclei from (**e**). Data represent mean; TOPBP1-negative: *n* = 101, TOPBP1-positive: *n* = 16 micronuclei.

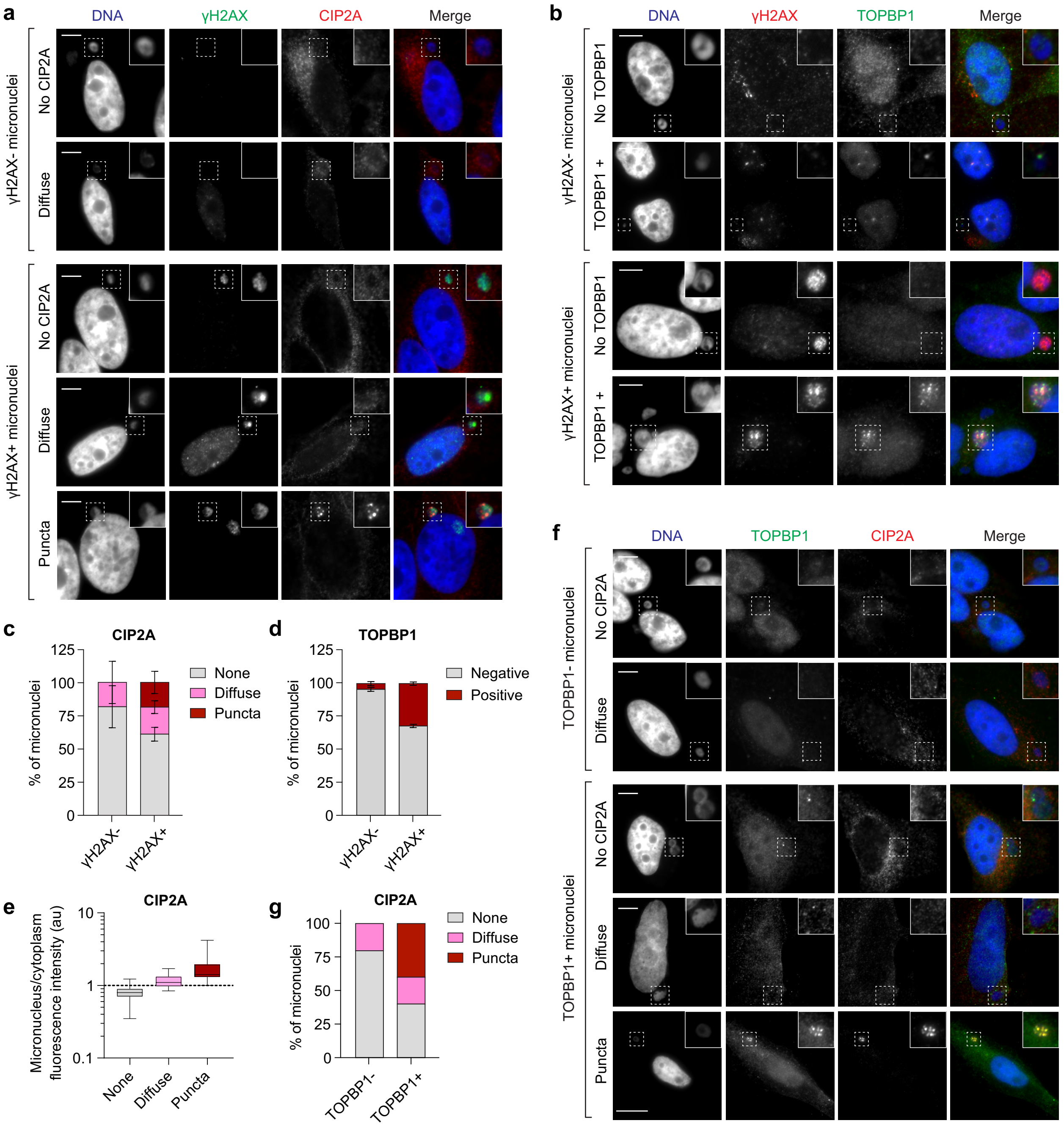


**Extended Data Figure 8. Interphase CIP2A-TOPBP1 patterns in ruptured micronuclei in HeLa cells. a-b)** Localization patterns of CIP2A (**a**) or TOPBP1 (**b**) in intact (γH2AX-negative) and ruptured (γH2AX-positive) micronuclei from HeLa cells treated with CENP-E and MPS1 inhibitors. Scale bar, 10 μm. **c)** Frequency of CIP2A localization patterns in intact (γH2AX-negative) and ruptured (γH2AX-positive) micronuclei. Data represent mean ± SEM; *n* = 3 independent experiments; γH2AX-negative: 119, γH2AX-positive = 76 micronuclei. **d)** Frequency of TOPBP1 localization patterns in intact (γH2AX-negative) and ruptured (γH2AX-positive) micronuclei. Data represent mean ± SEM; *n* = 3 independent experiments; γH2AX-negative = 179, γH2AX-positive = 51 micronuclei. **e)** Intensity measurements of the indicated CIP2A localization patterns in micronuclei compared to the cytoplasm. Box plot represents interquartile range with min-max; none: $n$ = 142, diffuse: $n$ = 39, puncta: $n$ = 13 micronuclei from 3 independent experiments. au, arbitrary units. **f)** Localization patterns of CIP2A and TOPBP1 in HeLa cells with micronuclei harboring random chromosomes. Scale bar, 10 μm. **g)** Frequency of CIP2A localization patterns in TOPBP1-negative and TOPBP1-positive micronuclei from (**f**). Data represent mean; TOPBP1-negative: $n$ = 84; TOPBP1-positive: $n$ = 9 micronuclei.

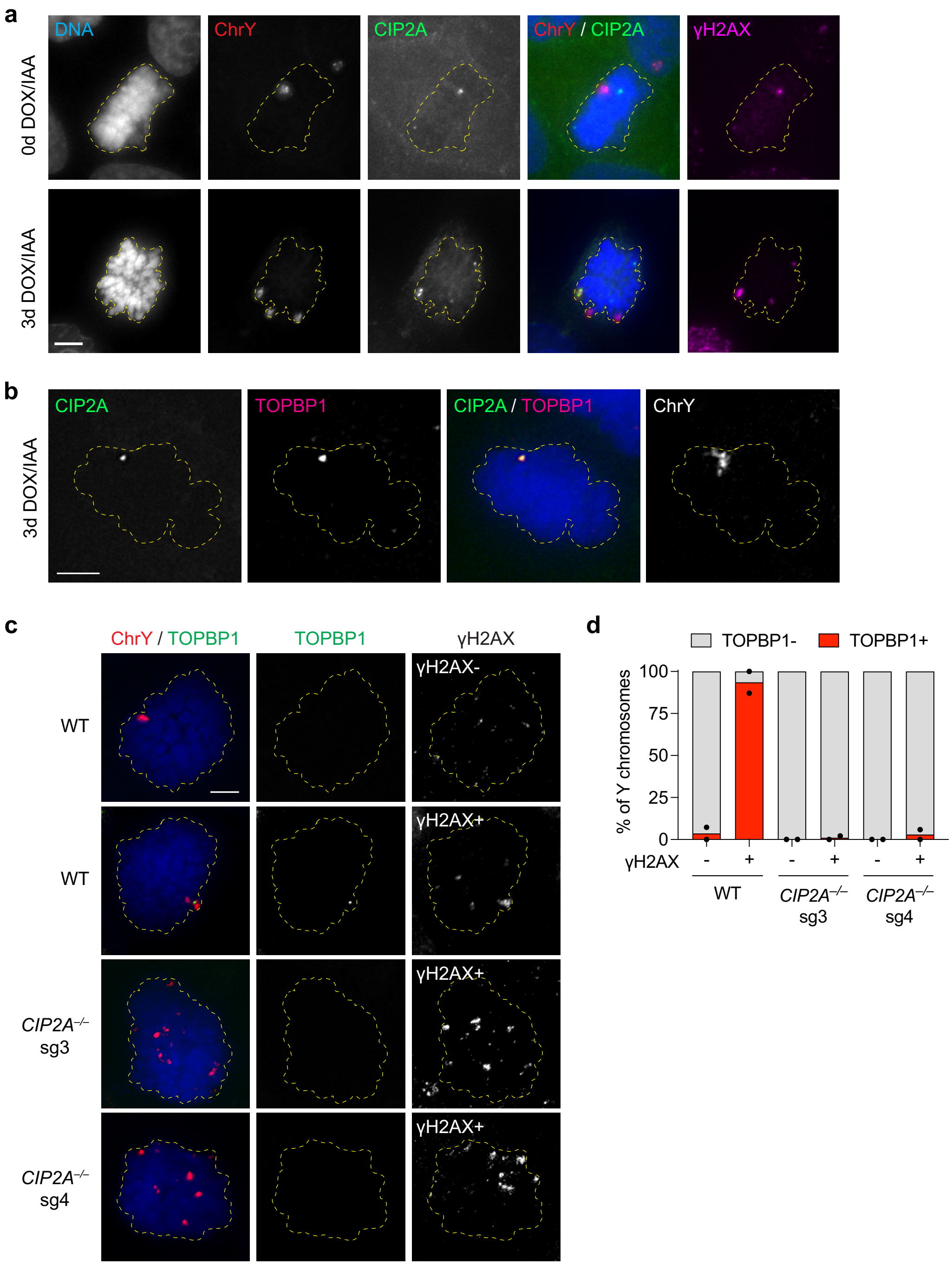


**Extended Data Figure 9. Mitotic localization of CIP2A and TOPBP1 on fragmented chromosomes.** **a)** Mitotic DLD-1 cells stained for CIP2A and H2AX and hybridized to chromosome paint probes. In untreated cells, CIP2A specifically co-localizes with spontaneous DNA lesions. Scale bar, 5 μm. **b)** Mitotic DLD-1 cells stained for CIP2A and TOPBP1 and hybridized to chromosome paint probes showing co-localization between CIP2A-TOPBP1 with the Y chromosome. Scale bar, 5 μm. **c)** Mitotic WT or CIP2A KO DLD-1 cells stained for TOPBP1 and H2AX and hybridized to chromosome paint probes. CIP2A loss prevents TOPBP1 recruitment to dispersed Y chromosome fragments. Scale bar, 5 μm. **d)** Quantification of (**c**). Data represent the mean of *n* = 2 independent experiments; left to right: 330, 114, 94, 124, 308, and 153 cells.

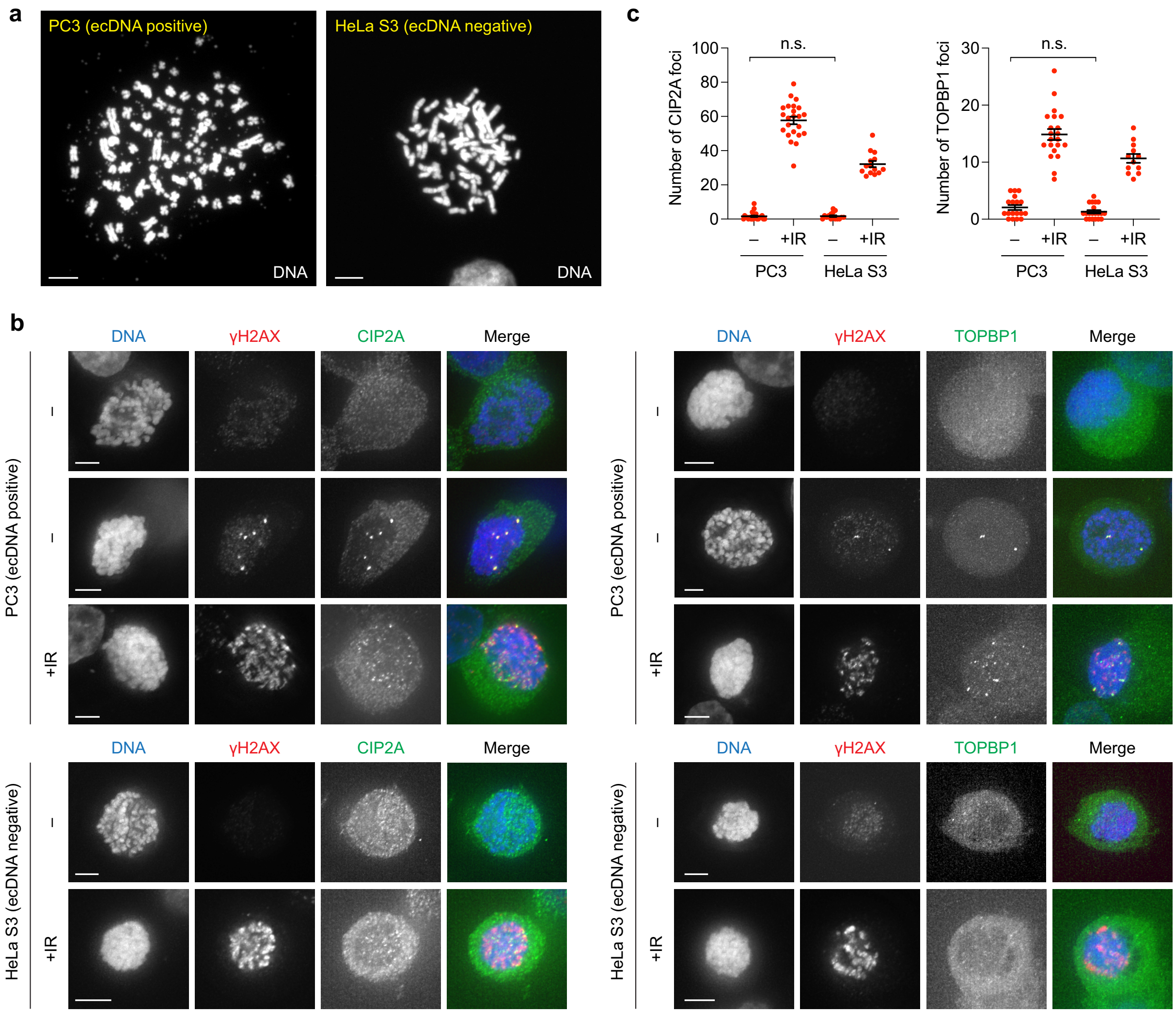


**Extended Data Figure 10. CIP2A-TOPBP1 does not associate with acentric extrachromosomal DNA (ecDNA) elements lacking DNA damage. a)** DAPI-stained metaphase spreads showing abundant ecDNAs in PC3 cells but not control HeLa S3 cells. Scale bar, 5 μm. **b)** CIP2A and TOPBP1 are not recruited to mitotic chromosomes in PC3 cells with ecDNAs in the absence of DNA damage. Examples of untreated and irradiated PC3 and HeLa cells arrested in mitosis and immunostained for CIP2A or TOPBP1. Scale bar, 5 μm. **c)** Quantification of CIP2A and TOPBP1 foci in (**b**). Data represent the mean ± SEM; $P$ = 0.8768 (ns) for CIP2A; from left to right, $n$ = 24, 23, 15, and 14 mitotic cells; $P$ = 0.1437 (ns) for TOPBP1; from left to right, $n$ = 18, 21, 20, and 12 mitotic cells; $P$-values calculated by two-tailed t-test comparing non-irradiated PC3 and HeLa S3 cells.

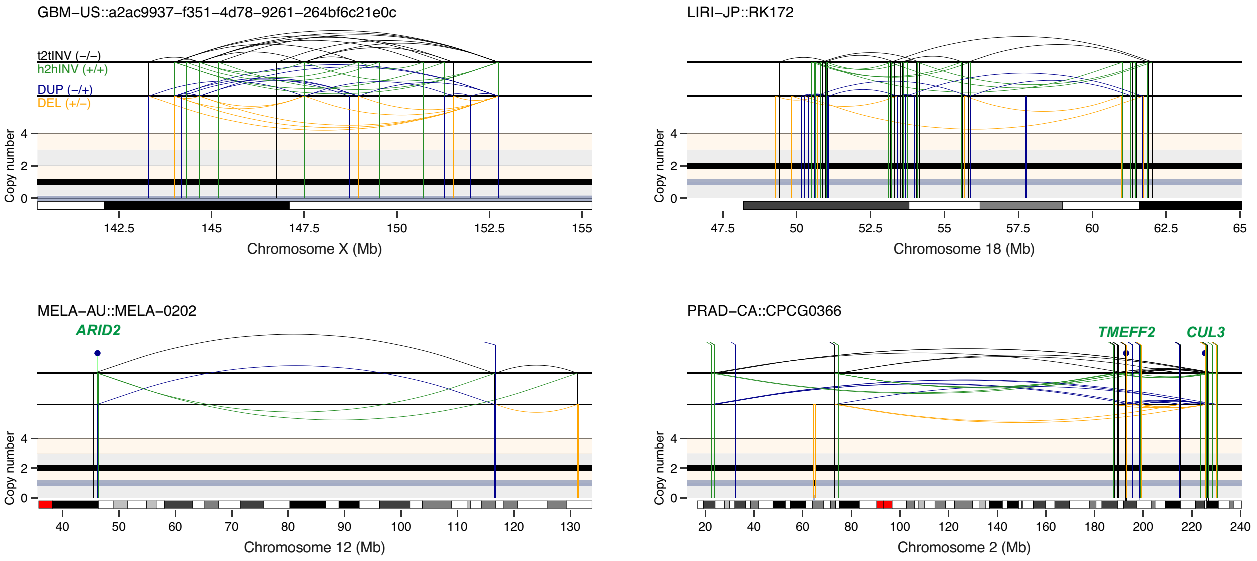


**Extended Data Figure 11. Additional examples of balanced chromothripsis events in cancer genomes.**

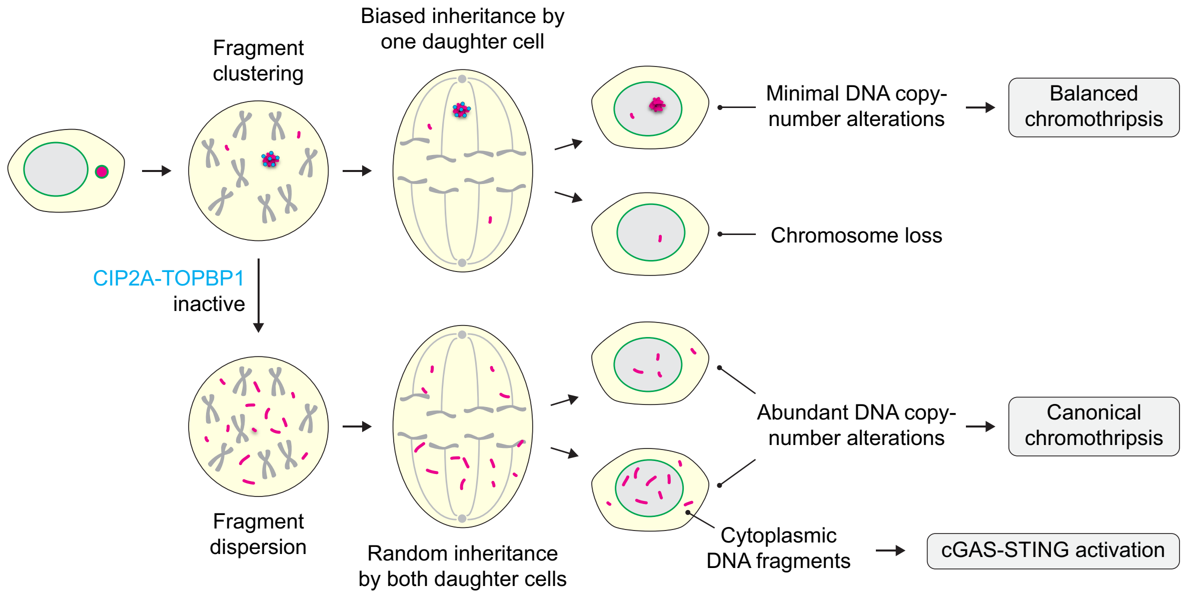


**Extended Data Figure 12. Graphical schematic depicting how mitotic clustering facilitates balanced and canonical rearrangements from chromothripsis.**

**Extended Data Table 1. List of balanced chromothripsis events disrupting cancer driver genes in the PCAWG cohort.**

**Supplementary Table 1. List of oligonucleotides used in this study.**
